# A memory-driven reinforcement learning model of phenotypic adaptation for anticipating therapeutic resistance in prostate cancer

**DOI:** 10.1101/2025.11.03.686133

**Authors:** Zahra S. Ghoreyshi, Shibjyoti Debnath, Pelumi D. Olawuni, Andrew J. Armstrong, Jason A. Somarelli, Jason T. George

**Affiliations:** Department of Biomedical Engineering, Texas A&M University, College Station, 77843, TX, USA; Translational Medical Sciences, Texas A&M University Science Center, Houston, 77030, TX, USA; Center for Theoretical Biological Physics, Rice University, Houston, 77005, TX, USA; Department of Medicine, Duke University, Durham, 27710, NC, USA; Duke Cancer Institute Center for Prostate and Urologic Cancers, Duke University Hospital, Durham, 27710, NC, USA; Department of Hematopoietic Biology and Malignancy, MD Anderson Cancer Center, Street, Houston, 77030, TX, USA

## Abstract

While contemporary cancer treatment strategies have significantly prolonged the lives of patients, therapeutic resistance remains a predominant cause of disease progression and cancerrelated deaths. Cancer therapy often induces gene regulatory responses that promote cell survival in the face of this therapy. Herein, we sought to develop a stochastic model of the response to repeat therapeutic challenge. This model integrates reinforcement learning to account for environmental history-dependent cellular transitions and growth dynamics. When applied to prostate cancer, this memory-driven adaptive model successfully captures the experimentally-observed dynamics of drug-sensitive and drug-resistant LNCaP cells under varying dosing schedules of androgen receptor blockade with enzalutamide (enza), significantly outperforming traditional transition models that lack history dependence. This performance is especially evident in the ability of our approach to robustly predict stochastic fluctuations in cancer cell population sizes across the entire disease trajectory, including subtle, later-emerging responses following initial therapy. The model was further evaluated by predicting the control of resistant cells in an enza environment by modeling inhibition of the p38/MAPK pro-survival stress axis, which was then validated experimentally. Lastly, we developed and applied a patient-calibrated model using prostate-specific antigen (PSA) data from clinical patient cohorts undergoing intermittent androgen deprivation therapy. Our model accurately predicts the PSA dynamics under repeated treatment cycles and effectively distinguishing between patients who respond and those who do not respond to treatment, thereby providing quantitative insight into prostate cancer progression. We anticipate that such adaptive modeling frameworks will be broadly useful for predicting cancer treatment outcomes and developing optimized adaptive therapeutic strategies tailored to patient-specific disease dynamics in additional cancer contexts.

## Introduction

Drug resistance remains a persistent obstacle to delivering durable treatment strategies in cancer therapy (1). Extensive prior work has established a myriad of pre-existing and acquired resistance mechanisms (2, 3), phenotypic plasticity and adaptation (4, 5) and cell state switching (6, 7), as well as complex cell–environmental mechanisms of resistance that emerge within cancer subpopulations (8, 9). Experimental and theoretical studies have framed cancer therapy resistance as an adaptive process, highlighting both genetic and non-genetic mechanisms of resistance that allow cancer populations to circumvent therapeutic targeting (10–27). Complementary mathematical and eco-evolutionary models of adaptive therapy have gone far to elucidate how treatment-induced selective pressures shape competition between sensitive and resistant subpopulations, wherein early therapeutic responses can provide predictive insight into patient-specific disease dynamics (28–31). Attempts to target the molecular pathways underlying resistance have yielded minimal improvements in patient outcomes (32, 33), reflecting the plasticity and multitude of mechanisms by which tumor cell adaptation can lead to therapeutic failure or evasion. Collectively, these studies demonstrate that cancer cells can enter reversible drug-tolerant states driven by non-genetic mechanisms, including chromatin remodeling (21), stochastic phenotypic switching (26), and the acquisition of rare transcriptional states (14–17). In this context, epigenetic adaptation serves to further facilitate drug resistance by inducing in cancer cell populations plasticity in phenotypic states. Building on these insights, recent theoretical advances suggest that the complex behaviors exhibited by cancer cells under treatment can be understood through the lens of decision theory. Specifically, cancer dynamics reflect an adaptive decision process, wherein cells balance trade-offs between proliferation and survival under fluctuating environmental pressures (23, 34, 35). This perspective not only provides a unifying framework for modeling phenotypic plasticity, but also enables the incorporation and evaluation of dynamic optimization in predictive models of cancer progression. The degree to which cancer cell dynamics can be explained by phenotypic switching using a memory-driven adaptive decision scheme is at present unknown.

To address these gaps, we develop and apply a general modeling framework that explicitly incorporates memory-driven phenotypic adaptation in response to treatment history. To evaluate the role of memory-driven adaptation, we first developed an integrated theoretical and experimental approach using repeat challenge of LNCaP (PCa) cell lines with standard-of-care androgen receptor pathway inhibitor, enzalutamide (enza). By estimating the growth rates of LNCaP cells exposed to periodically changing treatment environments, we observed fitness variations that align most closely with theoretical predictions of memory-driven adaptation in a regime that optimally balances adaptation speed and accuracy. Using an experimental system with dual fluorescence-based reporters to track high-resolution temporal dynamics over three weeks, we studied phenotypic adaptation in response to repeat drug exposures. Employing a reinforcement learning (RL) framework to model cell growth, death, and phenotypic transitions, we demonstrate that memory-driven adaptation can explain observed population-level phenotypic dynamics more accurately than models lacking memory mechanisms over a variety of treatment policies. We also demonstrate that incorporating memory-driven adaptation enables predictions of cell population behavior during inhibition of the p38/MAPK pro-survival (36) signaling pathway. We applied our model to study progression of clinical disease in the setting of longitudinal PSA monitoring from a trial of intermittent androgen deprivation therapy (ADT). Our model, when trained on initial dynamics for individual patients, accurately predicts long-term outcomes, successfully resolving patients that progress from those that exhibit long-term disease control based on their initial PSA time-course data. Collectively, our results demonstrate the utility of incorporating memory-driven adaptation into predictive models of cancer progression. We anticipate that this modeling approach will be broadly applicable to understanding therapeutic response and establishing prognostic markers of treatment resistance in other cancer contexts.

## Materials and Methods

To comprehensively investigate memory-driven phenotypic adaptation in PCa, we combined computational modeling and experimental validation into an integrated methodological framework. Experimentally, we conducted controlled in vitro studies using paired enza-sensitive and enza-resistant LNCaP PCa cell lines under cyclic and continuous drug exposures, as well as co-culture assays to assess cellular responses to dynamic drug switching. These experiments provided detailed data on growth dynamics, phenotypic adaptation, and the impact of therapeutic interventions. To complement these experimental analyses, we developed two distinct yet synergistic computational approaches: a discrete-time stochastic model to characterize population-level adaptation under periodically switching treatments, and a continuous-time reinforcement learning (RL) model to capture individual cell phenotypic transitions with high temporal resolution. The integration of these experimental and computational approaches allows us to validate model predictions rigorously and elucidate the critical role of memory-driven mechanisms in shaping cancer cell responses to fluctuating therapeutic pressures.

### Quantification of Growth Dynamics under Cyclic and Continuous Drug Exposure

To evaluate how cyclic and continuous exposure to enza influences phenotypic adaptation and growth dynamics, we conducted controlled *in vitro* experiments using paired enza-sensitive and enza-resistant LNCaP cells. LNCaP cells were obtained from The Duke University Cell Culture Facility. All cell lines were verified by STR profiling in conjunction with the Duke DNA Analysis Facility. Cells were tested weekly for mycoplasma contamination using the MycoAlert Mycoplasma Detection Kit (Lonza). Cells were maintained in RPMI supplemented with 10% fetal bovine serum and 1% penicillin-streptomycin and incubated at 37°C and 5% CO2. Paired enza-sensitive and -resistant LNCaP cells were derived by chronic exposure to increasing doses of DMSO or enza (36). A parallel enza-sensitive sub-line was treated with DMSO in equal volume to the enza-treated cells. Enza-sensitive and -resistant lines were labeled by transducing EGFP (enza-sensitive) or mCherry (enza-resistant) into each cell line as previously described (37).

To quantify cell growth dynamics during repeat drug challenge, LNCaP cells were seeded at a density of 2,000 cells per well in 48-well plates. After 24 hours, cells were treated with 20 µM enza or an equivalent volume of DMSO as a vehicle control for 24 hours. Following treatment, the drug-containing medium was removed and replaced with fresh drug-free medium. After an additional 48-hour recovery period, cells were subjected to either cyclic exposure to DMSO or enza, or continuous exposure to DMSO or enza for 24 hours. Cell confluence was monitored using IncuCyte S3 live-cell imaging (Sartorius) at 10X total magnification, with measurements taken every 24 hours over a period of 14 days.

### Phenotypic Response of Co-cultured Cells to Drug Switching

To quantify the response of sensitive and resistant cells to drug-switching, labeled LNCaP enza-sensitive and -resistant cells were plated as a co-culture at 1:1 ratio on 96-well plates at a seeding density of 1,000 cells/well. The next day, DMSO or enza (50 µM) was added, and confluence of enza-sensitive and -resistant populations was monitored by IncuCyte S3 live cell imaging (Sartorius). After one week, drug conditions were switched as indicated: DMSO-DMSO, DMSO-enza, DMSO-P38 inhibitor (SB203580), enza-enza, enza-DMSO, or enza-enza+P38 inhibitor (SB203580), to assess the influence of targeted inhibition on pro-survival phenotypes. Cellular responses and growth dynamics posttreatment switching were evaluated to elucidate the phenotypic adaptation of cancer cells to changing therapeutic pressures.

### Model Development of PCa Phenotype transition

Our mathematical modeling strategy integrates two complementary approaches to characterize PCa cell growth and adaptation under dynamic environmental conditions (Figure 1a). The first approach employs a discrete-time stochastic model of memory-driven adaptation to describe population dynamics in periodically switching environments (Figure 1b). The approach links the adaptation timescale to phenotypic memory of prior environmental conditions and is subsequently applied to experimental data from cells exposed to oscillating treatment regimens.

**Fig. 1.**
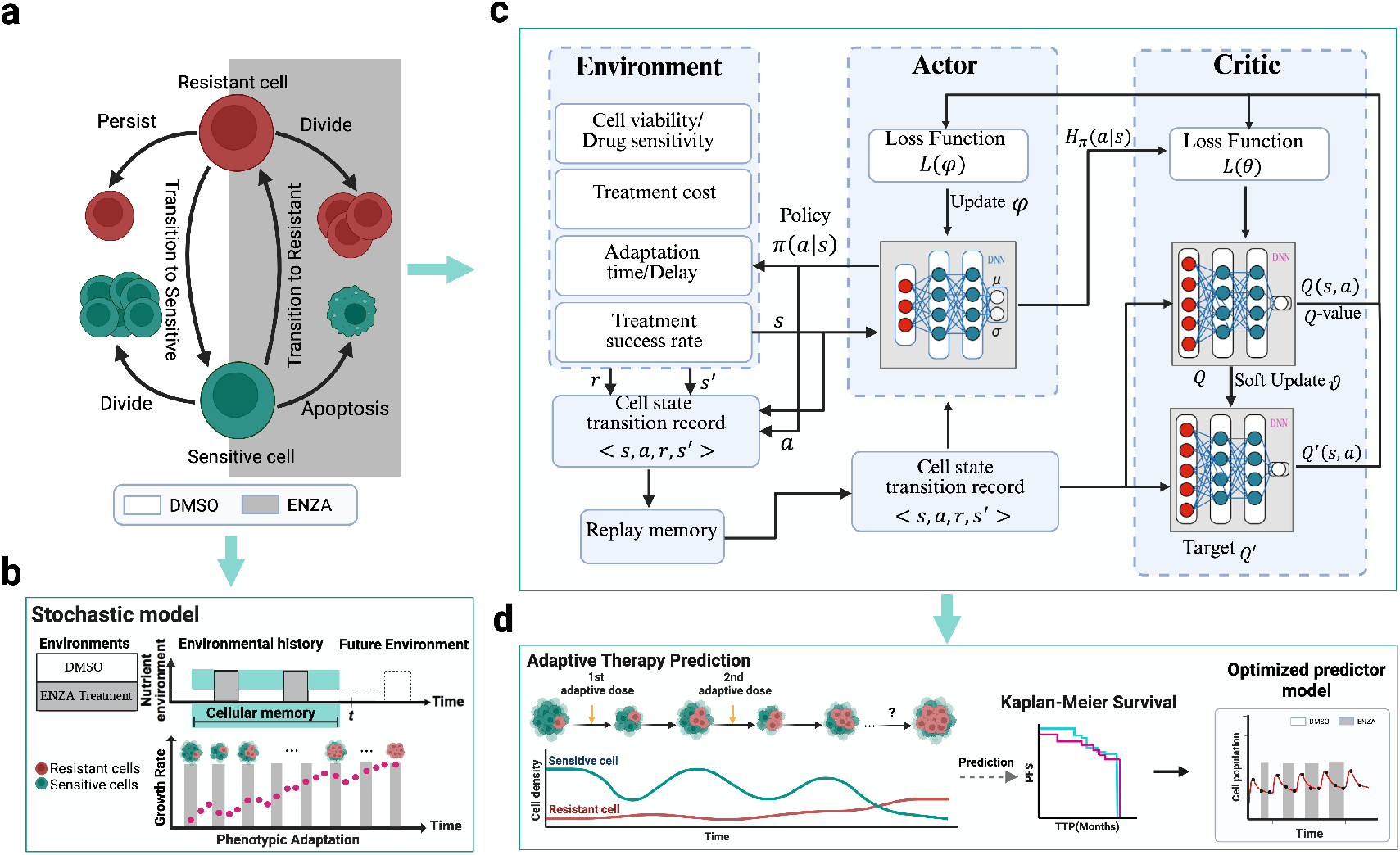
Framework overview. (a) Schematic representation of phenotypic subpopulations (enza-sensitive and enza-resistant) and their reversible transitions under different environmental conditions, including enza-treated and DMSO-control environments. This phenotype transition structure forms the basis for modeling adaptive dynamics. (b) Stochastic modeling framework used to identify the optimal memory window length that captures how cumulative environmental exposure influences phenotypic transitions. Reinforcement learning framework using the Soft Actor-Critic (SAC) algorithm to learn optimal treatment policies. The SAC model incorporates the environment-specific phenotypic adaptation dynamics and is trained on synthetic trajectories generated from the calibrated stochastic model. The learned policy outputs continuous treatment actions that adaptively respond to tumor states and memory-driven transitions. (d) Outputs of the RL model, including adaptive therapy predictions illustrating how dynamic treatment decisions influence sensitive and resistant cell populations over successive treatment cycles, and optimized predictor trajectories showing detailed cell population responses.

The second approach utilizes a continuous-time reinforcement learning (RL) model designed to describe the dynamics of individual cell phenotypes in experiments with high temporal resolution. In contrast to the discrete-time model, this RL framework explicitly accounts for heterogeneity in cell populations, allowing for detailed modeling of cell growth, death, and phenotypic transitions among distinct subpopulations Figure 1c. This model is designed to evaluate treatment efficacy and predict the effects of dynamic dosing schedules on population dynamics (Figure 1d).

#### Aggregate Population-Level Modeling

To estimate a general timescale on which a history of past environmental exposures supports experimental growth rates, we specialized a previously developed memory-driven stochastic model (34) to describe the dynamics of phenotypic adaptation in fluctuating environments. This model focuses on two distinct environmental states: DMSO and enza, in addition to two corresponding phenotypic states for PCa cells, a sensitive phenotype *S*_DMSO_ (optimized for growth in DMSO) and a resistant phenotype *S*_enza_ (optimized for growth in enza). This discrete-time model assumes that cancer cells can utilize a history of past environmental exposures to undergo phenotypic transition to the most beneficial state. Using this framework, growth rates for each phenotype can be represented at each time by:

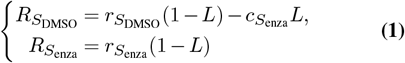

where *L*∈ { 0, 1 } represents the state of the environment with *L* = 0 (resp. *L* = 1) corresponding to a DMSO (resp. enza), environment, 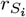 corresponds to the growth rates of the *S*_*i*_ phenotype in a DMSO environment (*L* = 0), and 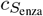 denotes a cost of the *S*_enza_ phenotype when enza is present.

To determine phenotypic switching, cells at period *n* in this model possess a history, or environmental ‘memory’, of the most recent *N* prior environmental states, in the form of an empirical average, *π*_*n*_. This average *π*_*n*_ is then dynamically compared to a critical value *π*_*crit*_ to determine the optimal phenotype. It was previously shown (34) that the size of *N* introduces a trade-off: larger memory sizes simultaneously improve the accuracy of environmental estimation and reduce the rate of adaptation to sudden shifts in the environment. Lower environmental values of *N* enable rapid adaptation, but lead to estimation errors and phenotypic mismatches.

This model is applied to aggregate population-level cell growth by determining per-cell growth rates for each phenotype through a two-pronged approach. For simulation data, growth rates were computed dynamically updating based on the environmental conditions at each time step using Eq. 1. Model-derived growth rates were compared to experimentally derived growth rates estimated by calculating the time derivative of cell population size.

#### Time-resolved Individual Phenotype Modeling

We developed a stochastic model to describe the growth dynamics of PCa cell populations under different treatment conditions. The model assumes a dominant phenotype at each time step and was calibrated using daily measurements obtained from a growth dynamics experiment. Phenotypic transitions and population growth rates were estimated based on lineage-traced experimental data in which enza-sensitive and -resistant populations were each fluorescently labeled with mCherry or EGFP. While suitable for lower-resolution time series, this modeling approach does not explicitly capture concurrent subpopulations or rapid transitions observed in higher temporal resolution settings, such as the phenotypic response experiment.

While the discrete-time stochastic model is appropriate for studying aggregate population dynamics, it lacks the resolution to capture the competing dynamics of sensitive and resistant cells observed in highly time-resolved data. To address this, we developed a deep reinforcement learning (DRL) framework, focusing on the Soft Actor-Critic (SAC) algorithm. The DRL approach functions as a robust *controller* in a closed-loop system, dynamically modulating treatment inputs in response to environmental feedback to influence phenotypic adaptation. The SAC introduces entropy regularization alongside reward maximization, ensuring a balanced exploration of the phenotypic landscape and also enables the model to capture the coexistence and transitions of multiple subpopulations over short timescales (see SI, sec. S1 for more details). Compared to the stochastic model, the SAC-based controller offers significant advantages in handling uncertainty, capturing phenotypic co-existence, and modeling rapid transition dynamics, although it requires increased computational effort.

#### Soft Actor Critic in PCa – Environment

Cancer adaptation complexity arises from phenotypic transitions driven by both environmental factors and subpopulation interactions, posing significant challenges for comprehensive modeling (38). To address the complexity of cancer adaptation driven by environmental factors and subpopulation interactions, we developed a systems control–based mathematical model that incorporates cell division, environment-specific selection, and history-dependent phenotypic transitions. This framework captures the essential dynamics shaping cancer population behavior, including selection, reversible phenotypic switching, and heritable modifications such as mutations and epigenetic changes (39, 40).

Central to our approach is the cumulative environmental memory window *I*(*t*), which encodes historical treatment exposure integrated over a defined period:

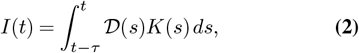

where *K*(*s*) is a kernel function capable of assigning greater weight to recent treatment exposures. This formulation allows the model to account for historical environmental conditions.

Using this environmental memory concept, we introduce the *phenotype transition matrix* Φ(*t*), which describes the transition probabilities between phenotypes as a function of the cumulative memory window and recent treatment history:

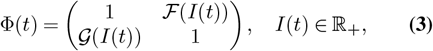

where ℱ (*I*(*t*)) and *G* (*I*(*t*)) are memory-driven functions explicitly defined as follows:

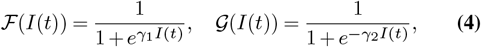

where *γ*_1_ and *γ*_2_ are learnable parameters that control the steepness and asymmetry of the transitions. These functional forms were convenient choices to reflect biologically relevant, nonlinear dynamics observed empirically. The logistic transformations ℱ (.) (resp. *G* (.)), governing transitions to the sensitive (resp. resistant) state, ensure smooth yet asymmetric switching between phenotypic states. As treatment pressure increases, ℱ (.) monotonically decreases the likelihood of transitioning to the sensitive phenotype, while *G* (.) correspondingly increases the likelihood of transitioning to the resistant phenotype (see SI Sec. S2 for more details).

Building upon previous work in modeling cancer adaptation (39–42), we integrate the phenotype transition matrix into a modified, time-dependent mixed-effect generalized Lotka–Volterra (tM-GLV) model based on the framework introduced in (43). This model captures population-level interactions between two primary phenotypic states: enza-sensitive (S) and enza-resistant (R) PCa cells. We assume that R-cells constitute a minority population but can continue to proliferate under conditions of androgen suppression. Environmental fluctuations further drive transitions between these phenotypes, reflecting the inherent heterogeneity of the tumor microenvironment. To enhance biological realism, the model includes two additional intermediate states, representing cells transitioning between sensitive and resistant phenotypes.

In our experimental setup, initially resistant cells are labeled red with a number (*X*_*RR*_) and initially sensitive cells are similarly labeled green (*X*_*SG*_). Following treatment initiation, these cells can undergo growth, death, or phenotypic transitions (*X*_*RR*_ to *X*_*SR*_ and *X*_*SG*_ to *X*_*RG*_) while retaining their original fluorescent labels (see SI sec. S1 for more details). This labeling method assumes monotonic growth behavior associated with the original phenotype, validated experimentally through chronic cultivation of resistant cells in enza for extended durations (>1 year), thereby ensuring consistency of phenotypic classifications throughout the experiment.

**Table 1.** Definitions of notations.

| Notation | Description | Units |
| --- | --- | --- |
| $X_G$ | $(x_{SG}, x_{RG})$, (sensitive, resistant) vector of cell counts (Green fluorescence) | 1* |
| $X_R$ | $(x_{SR}, x_{RR})$, (sensitive, resistant) vector of cell counts (Red fluorescence) | 1 |
| $R_G$ | $\text{diag}(r_{SG}, r_{RG})^\dagger$, inherent growth rate for two phenotypes | day<sup>-1</sup> |
| $R_R$ | $\text{diag}(r_{SR}, r_{RR})^\dagger$, inherent growth rate for two phenotypes | day<sup>-1</sup> |
| $\tilde{R}(D)$ | Effective treatment-induced cell death rate incorporating phenotype transition cost | day<sup>-1</sup> |
| $D$ | $\text{diag}(D, D)$, drug pressure: drug-induced decay, $D \in [0, 1]$ | 1 |
| $X$ | $\text{diag}(x_1, x_2)$, cell counts for two phenotypes | 1 |
| $\mathbf{1}$ | $(1, 1)$, vector of 1 | 1 |
| $\Phi(\cdot)$ | $[\mathcal{F}(I(t)), 1; 1, \mathcal{G}(I(t))]$, matrix of the phenotype transitions | 1 |
| $I(t)$ | memory window | 1 |
| $K$ | $\text{diag}(K_1, K_2)$, carrying capacity for two phenotypes | 1 |
| $\eta$ | Hyper-parameter in generalizing the phenotype transition matrix | 1 |

\* denotes no unit for this variable. †denotes a diagonal matrix with $r_1$ and $r_2$ as diagonal entries.

To model the observed phenotypic transitions, we developed a memory-driven, non-equilibrium framework governed by a system of differential equations. Unlike classical equilibrium-based models, this formulation captures time-dependent perturbations and maintains population dynamics within biologically feasible bounds (44, 45). The governing equations for initially enza-sensitive (green-labeled) and initially enza-resistant (red-labeled) subpopulations are:

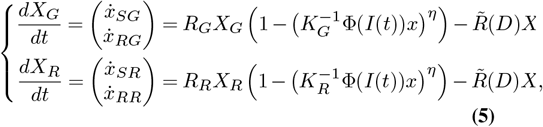

where *X*_*G*_ = (*x*_*SG*_, *x*_*RG*_)^⊺^ and *X*_*R*_ = (*x*_*SR*_, *x*_*RR*_)^⊺^ denote the cell subpopulation states labeled green (initially sensitive) and red (initially resistant), respectively. Parameters *R*_*G*_ and *R*_*R*_ represent phenotype-specific growth rates, while *K*_*G*_ and *K*_*R*_ correspond to the carrying capacities of these phenotypes. The phenotype transition matrix Φ(*I*(*t*)), previously defined (Eq. Eq. (3)), dynamically modulates phenotypic transitions based on environmental memory *I*(*t*). Parameter *η* controls the strength of density dependence, while the term 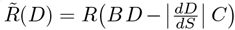 captures the effective rate of treatment-induced cell death, where *D* denotes the administered drug intensity, *B* = diag(*β*_1_, *β*_2_) collects the phenotype-specific drug–effect coefficients (*β*_1_ for sensitive cells and *β*_2_ for resistant cells), and 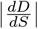 reflects the magnitude of treatment change across phenotypic states, with *S* representing the phenotypic state variable. The derivative 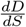 is used to detect transitions in treatment, either from no drug to drug exposure or vice versa, and its absolute value ensures that the transition cost is applied regardless of direction. The matrix *C* = diag {*C*_1_, *C*_2_ } encodes phenotype-specific costs associated with such transitions. When no change in treatment occurs, 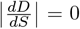, and no penalty is applied. This formulation ensures that adaptation to fluctuating treatment regimes incorporates a biologically-meaningful cost associated with phenotypic switching.

### Model-informed treatment planning with reinforcement learning

In closed-loop control systems, periodic feedback reduces dependency on precise trajectory knowledge by approximating uncertainty and stochasticity, enabling predictive dynamical tracking despite incomplete mechanistic understanding of cancer adaptation (46). This feedback-driven approach ensures robustness against dynamic changes in tumor behavior and parameter fluctuations. In our study, we implement reinforcement learning (RL) as the closed-loop controller, leveraging distributed modeling (Figure 2) to explicitly account for intratumoral heterogeneity and cellular variability. By considering multiple interacting cellular states and diverse tumor subpopulations, this distributed RL approach can effectively capture and respond to tumor evolution in real-time.

**Fig. 2.**
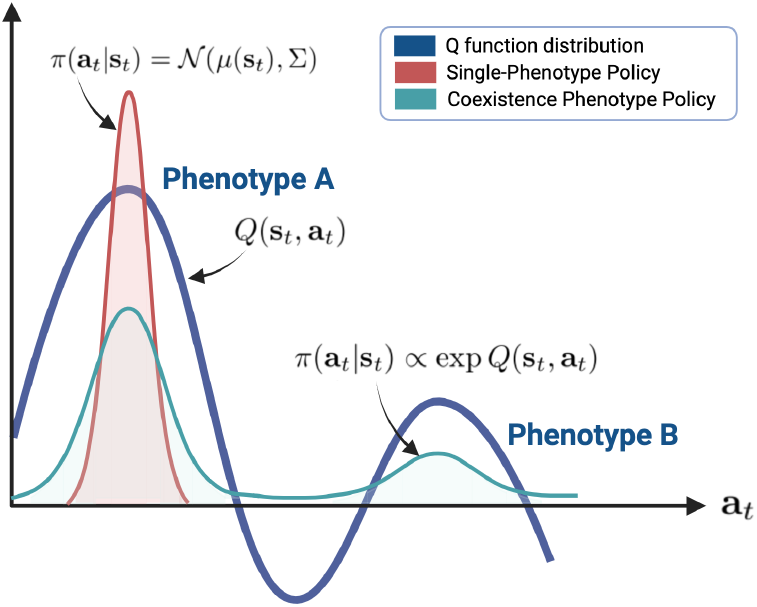
Visualization of the SAC policy and action distribution in continuous space. The red distribution represents the SAC policy for a single phenotype, modeled as a Gaussian *π*(*a* _*t*_|*s*_*t*_) = *N* (*µ*(*s*_*t*_), Σ), centered on the peak of the Q-function *Q*(*s*_*t*_, *a*_*t*_). The green distribution shows how the SAC policy adapts to incorporate Q-function values for the coexistence of phenotypes, enabling exploration and exploitation with *π*(*a*_*t*_|*s*_*t*_) ∝ exp(*Q*(*s*_*t*_, *a*_*t*_)).

Modern RL algorithms are broadly categorized into value-based and policy-based methods (47). Policybased algorithms such as Deep Deterministic Policy Gradient (DDPG) (48), Trust Region Policy Optimization (TRPO) (49), Proximal Policy Optimization (PPO) (43), and Soft Actor-Critic (SAC) (50) have become increasingly popular. Each algorithm exhibits distinct advantages and challenges. DDPG, an off-policy actor-critic method, performs effectively in continuous action-state spaces but is sensitive to hyperparameter tuning. TRPO uses Kullback-Leibler (KL) divergence (51) constraints to stabilize policy updates, yet it incurs computational overhead due to second-order optimization. PPO provides stable training and ease of implementation, suitable across action-space scenarios. Among these, SAC stands out due to its entropy-regularization mechanism, which encourages exploratory behavior and robustly captures diverse tumor phenotypic dynamics. Given our need to model heterogeneity and adaptively respond to patientspecific tumor evolution, SAC’s capacity for capturing complex stochastic behaviors makes it particularly suited for our approach. Nonetheless, SAC required careful hyperparameter optimization tailored to each patient scenario to ensure stability and avoid divergence, as demonstrated by our empirical results.

#### Algorithm 1

Adaptive Treatment using SAC — Training ODE and SAC models

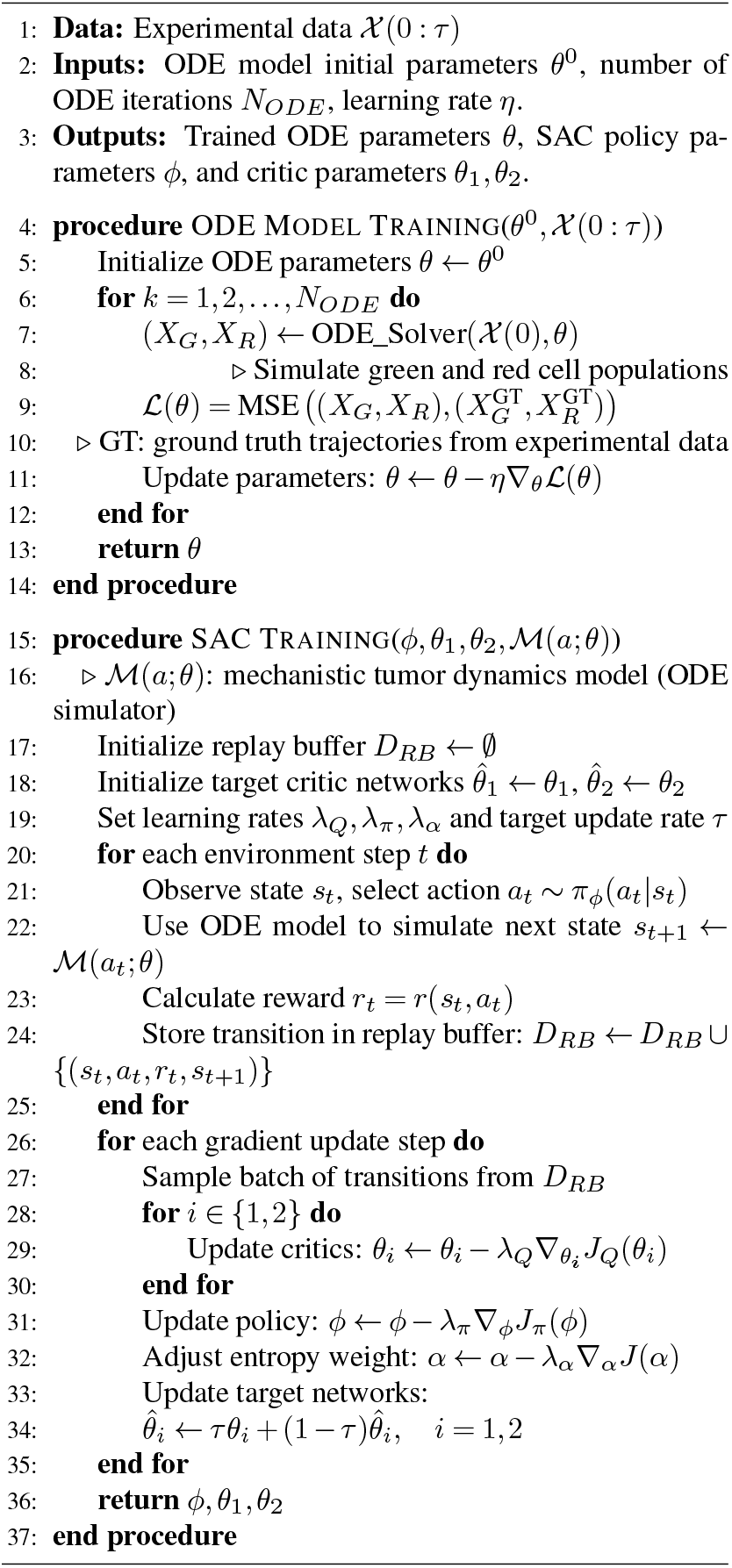

#### Algorithm 2

Adaptive Treatment using SAC — Adaptive Therapy Deployment

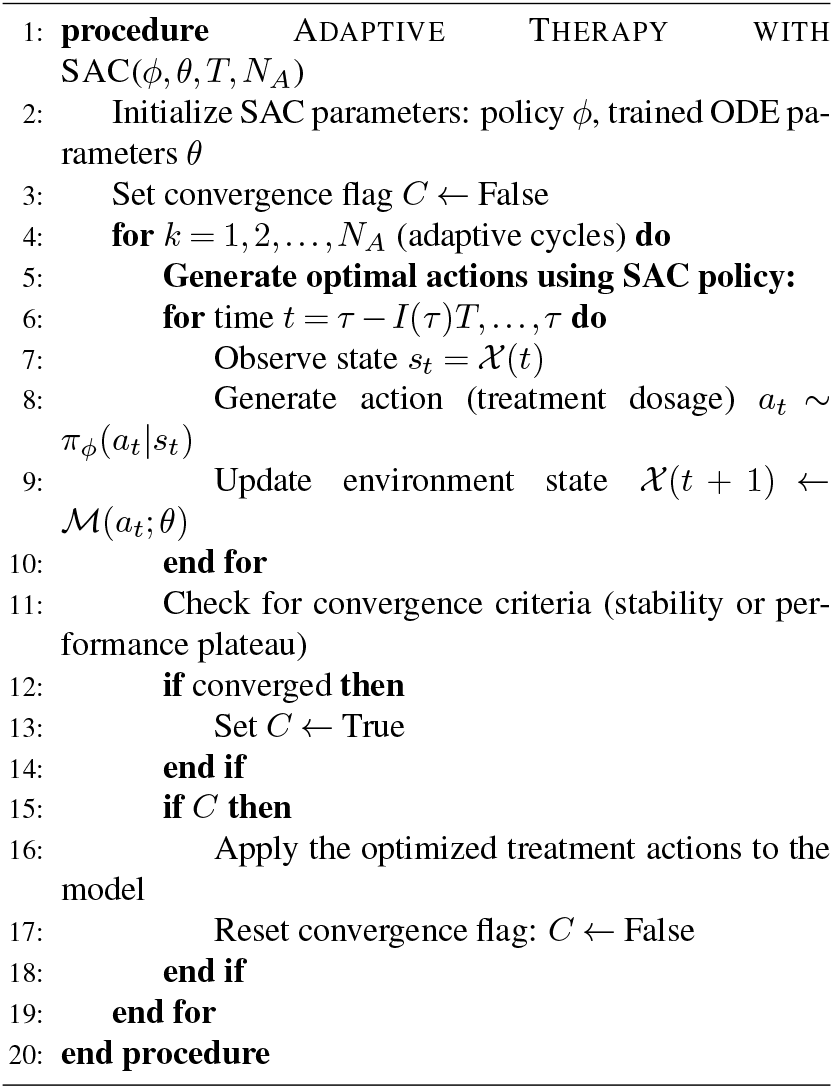

The SAC algorithm is designed to jointly optimize three core objectives: 1) learning accurate state-action value estimates (critic), 2) refining the policy (actor) to maximize long-term reward, and 3) dynamically balancing exploration and exploitation through entropy regulation. A full summary of the SAC training process, including ODE parameter optimization and policy learning, is provided in Algorithm 1. Each objective has a dedicated loss function, contributing to robust and stable policy learning in continuous action spaces. The critic network is trained to minimize the temporal difference (TD) error, defined as the difference between the predicted Q-values (*Q*-functions) and target Q-values. The Q-value *Q*_*θ*_(*s*_*t*_, *a*_*t*_) represents the expected cumulative future reward when taking action *a*_*t*_ in state *s*_*t*_, following policy *π* thereafter. Specifically, SAC uses two critic networks, parameterized by *θ*_1_ and *θ*_2_, to mitigate positive bias in Q-value estimation. Once training is complete, Algorithm 2 outlines how the optimized SAC policy is deployed in an adaptive therapy setting to generate patient-specific treatment schedules.

Training stability is further enhanced by a *replay buffer*, which is a memory mechanism that stores experiences (state transitions, actions, rewards) from previous interactions. At each training step, batches of these experiences are randomly sampled from the replay buffer, reducing correlations between sequential data and stabilizing learning. The critic loss for each Q-function (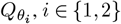) is given by:

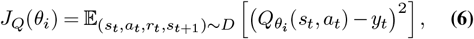

where the expectation is taken over a batch of experiences sampled from the replay buffer *D*, which contains previously observed transitions (*s*_*t*_, *a*_*t*_, *r*_*t*_, *s*_*t*+1_). The target value *y*_*t*_ is computed as:

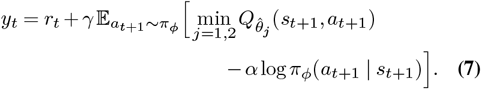

Here, *r*_*t*_ denotes the immediate reward obtained from transitioning from state *s*_*t*_ to *s*_*t*+1_ after taking action *a*_*t*_, and *γ* ∈ [0, 1) is the discount factor accounting for time value of future rewards. The term *α* log *π*_*ϕ*_(*a*_*t*+1_ | *s*_*t*+1_) introduces entropy regularization, thereby promoting stochastic exploration and encouraging diverse action selection.

To improve the policy, the actor network, parameterized by *ϕ*, is optimized to maximize expected returns while explicitly promoting exploration via entropy maximization. The policy loss function is defined as:

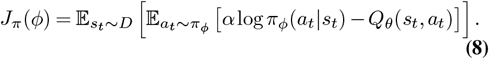

Minimization of this loss encourages actions with both high expected cumulative reward (high *Q*_*θ*_(*s*_*t*_, *a*_*t*_)) and high entropy (high uncertainty or variability), thereby preventing premature convergence to deterministic, potentially suboptimal actions.

In parallel, SAC incorporates an adaptive entropy temperature parameter *α*, which dynamically adjusts the entropy’s influence during training to maintain an optimal balance between exploration and exploitation. The temperature parameter is optimized by minimizing:

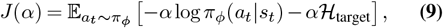

where ℋ_target_ is a predefined target entropy determining the desired policy stochasticity. By automatically adjusting *α*, the algorithm maintains sufficient exploration throughout the training process. Combining these components results in the complete SAC objective function, which the policy seeks to maximize:

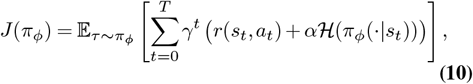

where *τ* represents a trajectory sampled according to the current policy *π*_*ϕ*_, and ℋ (*π*_*ϕ*_(| *s*_*t*_)) denotes the entropy of the policy at state *s*_*t*_. This entropy-augmented formulation allows for effective exploration of the continuous action space, particularly beneficial in adaptive treatment planning scenarios where multiple phenotypic states coexist and evolve stochastically. Figure 2 illustrates this concept, showing how SAC policy distributions adapt to diverse phenotypic contexts. The red Gaussian distribution exemplifies a narrowly optimized policy suited for a single phenotype scenario, while the broader green distribution illustrates adaptive stochasticity, highlighting the policy’s flexibility to support phenotype coexistence and effectively balance exploration with exploitation across treatment cycles.

### Clinical dataset

We analyzed data from the prospective Canadian Phase II trial conducted by Bruchovsky *et al*. (2006), which evaluated intermittent ADT in 109 men with non-metastatic, hormone-sensitive prostate cancer who exhibited biochemical recurrence (rising PSA) after radiotherapy (52). In this study, all patients first underwent a 36-week induction phase of continuous androgen suppression using cyproterone acetate and leuprolide acetate. Only those whose PSA levels fell below 4.0 *µ*g/L at both week 24 (≈168 days) and week 32 (≈224 days), with a stable or decreasing slope, were defined as responders and were eligible to enter the intermittent phase. Patients who failed to meet this induction criterion were classified as non-responders and were excluded from further follow-up. A total of 72 patients met criteria for the intermittent phase.

During the intermittent phase, treatment was suspended when PSA dropped below 4.0 *µ*g/L and resumed when PSA rose to 10 *µ*g/L. “Progression to androgen-independent disease” was defined as three consecutive PSA increases above 4.0 *µ*g/L despite castrate levels of serum testosterone The authors reported a mean time to progression of approximately 165 weeks. Thus, all patients represented in the publicly available intermittent-therapy dataset were responders by design and were later separated into three outcome groups: those who experienced biochemical progression as defined above (*n* = 12), those who developed radiographically confirmed metastasis during follow-up (*n* = 5), and those who remained progression-free over the follow-up period (*n* = 55).

### Optimal Dosing Strategies Derived from RL Policy

To initialize our framework, we first pre-trained the network on a small subset of patients, which provided a reliable starting point for individualized calibration. For each new patient, calibration was then performed using only the earliest 10–15% of available PSA timepoints. This calibration window was selected because it typically spans both phases of the initial treatment response trajectory: an early PSA decline following therapy onset and the subsequent rebound. Once calibrated, the patient-specific model environment was coupled with a model-free Soft Actor–Critic (SAC) agent. This setup allowed us to evaluate the predictive reliability of the memory-driven model and to derive optimized dosing policies tailored to each patient’s tumor dynamics.

Our implementation utilizes the time-dependent mixed-effect generalized Lotka–Volterra (tM-GLV) model to simulate cellular dynamics and train the RL agent for optimal drug scheduling. At each time step, microenvironmental data—including cell population counts and the ratio of enza-resistant (enza-R) to enza-sensitive (enza-S) cells—serve as state inputs. The SAC agent iteratively refines its policies by interacting with this simulated environment. The value function integrates therapeutic effectiveness, phenotype transitions, and cumulative drug dosage penalties to balance treatment efficacy against potential side effects (see SI sec. S3 for more details).

A primary objective of our adaptive therapy framework is to maintain a minimal yet stable population of resistant cells. While reducing overall drug dosage can help suppress resistance, it may also allow sensitive cell populations to proliferate unchecked, potentially resulting in a critical tumor burden. In light of this issue, we incorporate a progression-free survival component into the step reward function. This addition incentivizes the reinforcement learning agent to dynamically maintain a balanced cell population over time.

## Results

Below we detail the main findings of our modeling and experimental approach. Full details may be found in SI and Methods sections.

### Cell Growth in Periodic Environments Strikes an Optimal Memory Trade-Off between Adaptation Speed and Accuracy

Phenotypic adaptation to temporally fluctuating drug conditions presents a fundamental challenge in cancer treatment. To address whether PCa cells exhibit memory-driven transitions that dynamically enhance fitness under periodic drug exposure, we specialized and applied a previously established theoretical model of memory-driven adaptation (34). This conceptual model frames phenotypic switching (e.g., between drug-sensitive and drug-tolerant states) as a cellular “decision” influenced by a memory of past environmental exposures. This Bayesian inference-based model relates phenotypic transition rates across treatment cycles to a defined memory capacity, allowing quantitative predictions of adaptive fitness under dynamic treatment conditions. The model explicitly assumes that prior environmental states influence cellular phenotype so that growth rates dynamically improve over time in an oscillating environment, in contrast with an adaptation scheme lacking memory that would generate cyclical fitness behavior.

To experimentally test these predictions, we conducted experiments using the enzalutamide (enza)-sensitive LNCaP PCa cell line subjected to periodically fluctuating drug conditions. Cells were cultured in 48-well plates and exposed to alternating 24-hour intervals of low-dose enza and drug-free media over a two-week period (Figure 3a). Total cell confluence was monitored every 24 hours via live-cell imaging, enabling non-invasive tracking of population dynamics. Per-cell growth rates were subsequently derived by calculating the difference quotient of the population size during exponential growth phases. Experimental results demonstrated i) an initial decrease in cell proliferation rates upon initial enza exposure cycles, followed by ii) a gradual stabilization and iii) subsequent increase in growth rates with continued cycling (Figure 3b,c), which aligns qualitatively with with theoretical predictions of memory-driven phenotypic adaptationunder oscillating environments.

**Fig. 3.**
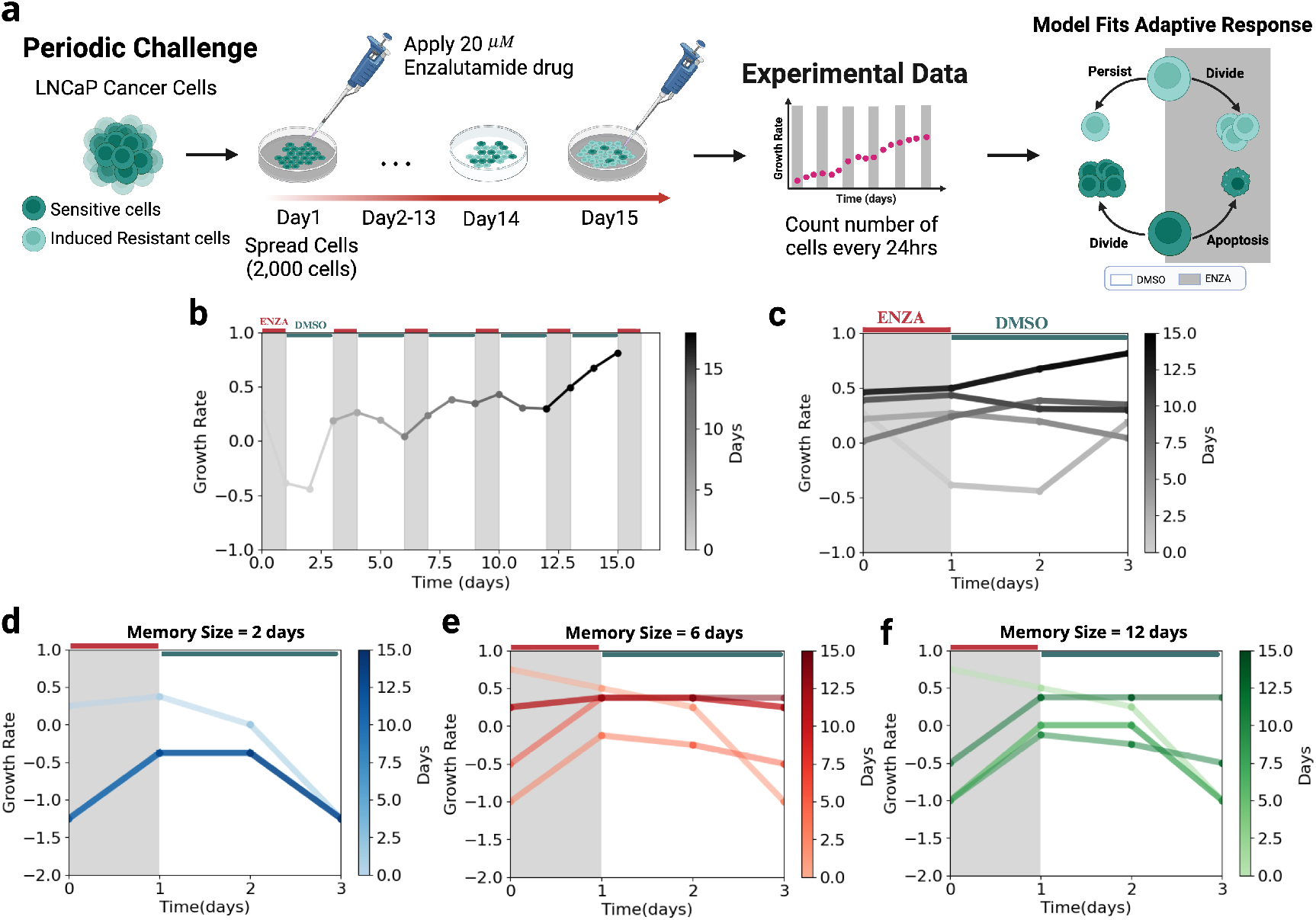
(a) Experimental approach to studying induced phenotypic resistance under periodic challenge: Time-series data collected over 24 days with 24-hour resolution, tracking overall PCa cell population dynamics under cyclic and continuous enza treatment. This experiment was used to calibrate the stochastic model and identify the optimal memory window size for modeling phenotypic adaptation. The simplified phenotype transition framework in this setting includes only growth, death, and persistence within the same state. (b) Experimental growth dynamics of enza-sensitive and enza-resistant LNCaP PCa cells collected with a treatment cycle of 3 days. The gray-to-black color gradient represents different experimental days, with lighter gray indicating earlier measurements and darker gray indicating later measurements. (c) Plot of the same experimental data in (b), with each line showing the short-term growth rate trajectory for a given day, allowing direct comparison across days under enza and DMSO conditions. Model predictions using a memory window of 2 days, which capture the adaptive response but do not fully reflect the temporal dynamics observed experimentally. (e) Model predictions using a 6-day memory window, which yield the optimal balance between convergence speed and fit quality, as indicated by the lowest mean squared error. Model predictions using a memory window of 12 days, where slower convergence results in a higher mean squared error relative to the 6-day window.

To quantitatively evaluate this theoretical prediction, we fixed the growth rates of enza-sensitive and induced enza-resistant cells, and we systematically varied the memory capacity as a single fitting parameter within our computational model. The experimentally derived growth rates as a function of period number (Figure 3c) were subsequently compared to model-driven predictions (Figure 3d-f). Using this approach, we find that the computational model fit the data best when using a memory capacity of six days, corresponding to twice the experimental drug treatment cycle (Figure 3e).

Intriguingly, when comparing the family of adaptation models having variable memory capacity, it turns out that the memory capacity most closely matching the experimental observations is optimal in the sense that it maximizes aggregate cellular fitness over the 15-day experimental window. The experimental dynamics were consistent with an intermediate memory response of 6 days. In contrast, our model predicted that shorter memory windows (e.g., two days; Figure 3d) were poorly adapted to the oscillating environment and exhibited low growth rates, reflective of insufficient and inaccurate environmental estimation. Conversely, larger memory responses (e.g., twelve days; Figure 3f) were predicted to provide minimal improvement in the accuracy of environmental estimation over the intermediate-memory case, but instead exhibited much slower dynamics to settle on the optimal phenotype, thereby reducing total aggregate fitness. Given our initial results supporting a memory-driven, optimally adaptive process, we proceeded to further explore the dynamics of phenotypic adaptation by integrating highly time-resolved experimental observations with a specialized dynamic programming modeling approach.

### Memory-Driven Phenotypic Adaptation Captures the Growth Dynamics of Highly Time-Resolved Phenotypic Subpopulations

The above observations at the bulk population level prompted us to investigate in more detail how dynamic environmental exposures might differentially influence enza-sensitive and enza-resistant subpopulations. We hypothesized that similar memory-dependent adaptation dynamics could manifest within a heterogeneous population of cancer cells exhibiting differential enza sensitivity. To mathematically describe these dynamics, we extended our memory-driven adaptation model by incorporating distinct enza-sensitive (enza-S) and enza-resistant (enza-R) cell populations, allowing each to exhibit phenotype-specific responses to prior environmental states. The refined framework explicitly models gradual, memory-dependent transitions between sensitive and resistant phenotypes, characterized by intermediate states and adaptive growth dynamics contingent upon the cells’ historical exposures. We hypothesized that variations in environmental memory would critically shape the subpopulation growth trajectories, influencing how rapidly each phenotype adapts to fluctuating drug conditions.

We validated this theoretical framework experimentally through high-resolution time-series analyses. Enza-sensitive (enzaR; EGFP-labeled) and enza-resistant (enzaS; mCherry-labeled) LNCaP cells were co-cultured at a 1:1 ratio in 96well plates and subjected to clearly-defined, periodic drug regimens. Initially, cells were exposed either to DMSO or enza (50 µM) for seven days, followed by continued exposure to either the same initial condition or an alternative condition for an additional seven days. Fluorescently labeled subpopulations were monitored hourly using IncuCyte live-cell imaging (Figure 4a). Our memory-driven adaptation framework was then fitted to the experimental data.

**Fig. 4.**
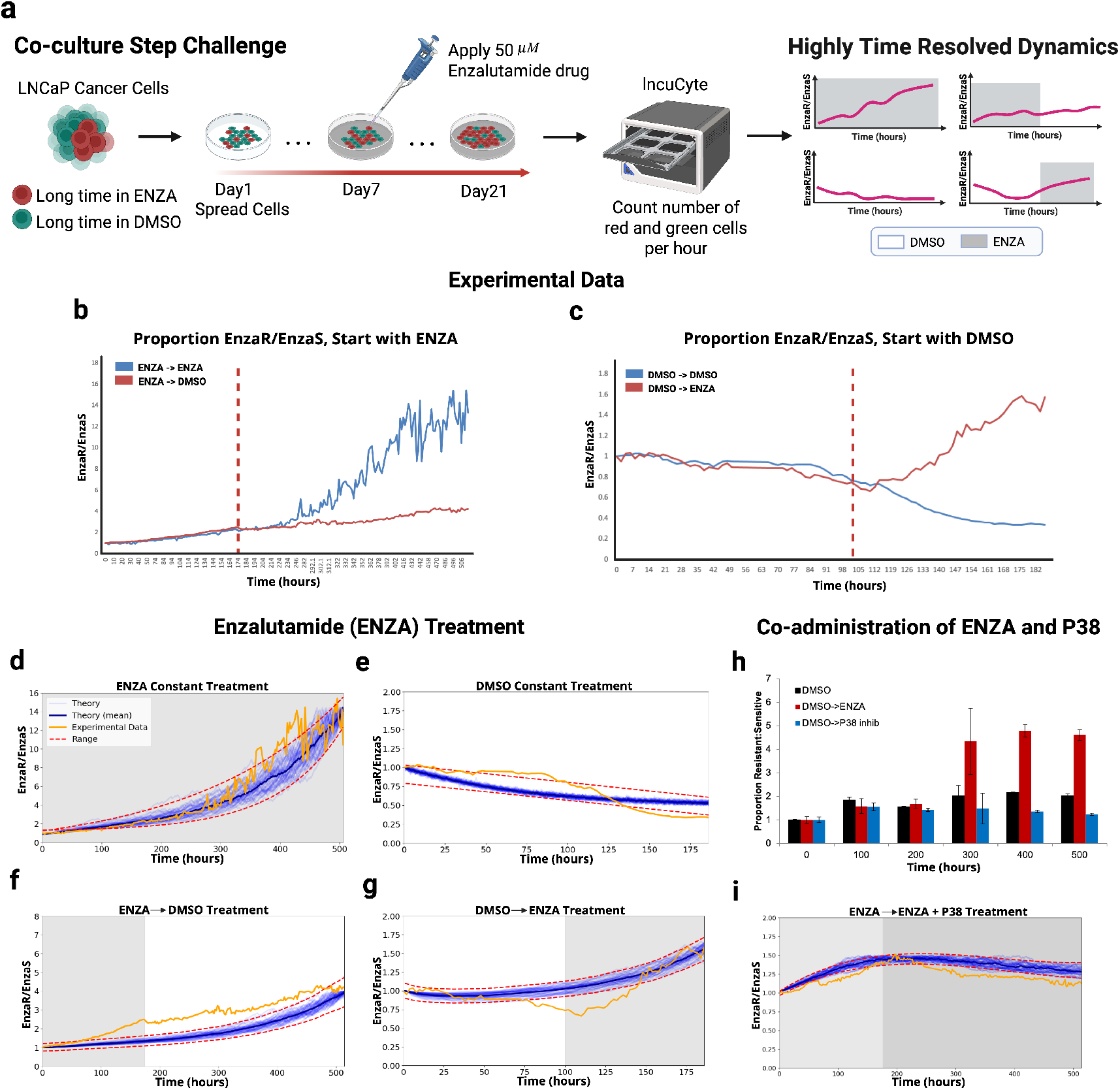
Population dynamics of enza-sensitive and enza-resistant co-culture under variable drug environments (a) High-resolution (hourly) imaging data from co-cultured enza-sensitive (EGFP-labeled) and enza-resistant (mCherry-labeled) LNCaP cells treated with enza, DMSO, or P38 inhibitors. This experiment was used to train the full phenotype transition model and identify optimal treatment strategies. (b) Experimental time-series of the proportion of enza-resistant to enza-sensitive cells (enza-R/enza-S) starting with enza treatment. (c) Experimental time-series of the proportion of enza-resistant to enza-sensitive cells starting with DMSO treatment. (d–I) Time-resolved growth dynamics of enza-sensitive (green) and enza-resistant (red) PCa cells under various treatment regimens. Panel (d) shows cells receiving two consecutive cycles of enza treatment; panel (e) depicts cells treated with two consecutive cycles of DMSO; panel (f) illustrates a regimen in which treatment begins with enza followed by DMSO; panel displays a regimen beginning with DMSO followed by enza; and panel (h,I) presents the dynamics under co-administration of DMSO, enza and a P38 inhibitor. Data were obtained from 1:1 co-cultures of LNCaP cells monitored in real time using live-cell imaging. These results were used to optimize the memory-driven adaptive model and to assess the distinct adaptive responses elicited by each treatment protocol.

Using this approach, we found that our computational model was able to explain the growth dynamics of enza-sensitive and enza-resistant mixtures under a variety of environmental perturbations with the ratio enzaR/enzaS reported for each case (Figure 4b-i). As expected, treatment with enza followed by DMSO results in lower ratios relative to constant enza exposure due to the growth of the enzaS population (Figure 4c). Similarly, treatment with DMSO followed by enza results in the enrichment of the enzaR population relative to constant exposure to DMSO, where the enzaS cells grow more rapidly (Figure 4d). To establish the added utility of incorporating memory-dependent adaptation, we compared the ability of the memory-driven adaptation model to explain the experimental data against that of a simpler ‘null’ model lacking historical environmental memory (see SI sec. S4 for details). Both models were independently fit to the experimental data with the stringent requirement of simultaneously explaining constant or switching enza/DMSO environments, and optimal parameters for each were determined based on best-fit criteria. We found that memory-driven model predictions significantly outperformed the null model in explaining experimentally-observed responses to concurrent environments (Figure 4e-g). These results support the value of incorporating cellular memory in describing empirically-observed adaptation dynamics, suggesting that the incorporation of environmental history in models of cellular decision-making may provide important insights into adaptive behavior in phenotypically-heterogeneous cell populations. Given our model’s initial ability to capture the above empirical dynamics, we next proceeded to evaluate our model’s ability to predict the dynamics under additional therapeutic strategies.

#### Predicting Growth Stasis with Combined Enzalutamide and Timed P38 Inhibition

Given that our calibrated model accurately reproduced phenotypic dynamics under enzalutamide therapy, we next sought to evaluate its predictive capacity in combination treatment scenarios aimed at suppressing the outgrowth of enza-resistant subpopulations. In particular, we focused on the P38 MAPK pathway, which is known to mediate stress-response signaling and support adaptive survival in enza-resistant prostate cancer cells (36). Based on this mechanistic role, we hypothesized that targeted inhibition of P38 would antagonize memory-driven resistance by selectively impairing the survival of resistant subpopulations. We specifically examined two clinically-relevant strategies: (1) whether P38 inhibition during enza treatment could shift the phenotypic balance in favor of sensitive cells, and (2) whether co-administration of enza and a P38 inhibitor could durably suppress both sensitive and resistant subpopulations, thereby delaying overall tumor progression.

To explicitly incorporate the effects of the P38 inhibitor into our modeling framework, we introduced a binary indicator variable, *D*_inhibitor_, to represent the presence or absence of inhibitor treatment:

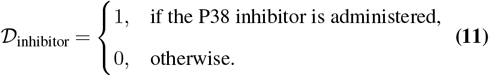

We then modified the previously defined phenotype transition matrix (Φ(*t*)) to include the inhibitor effects explicitly, yielding the following updated formulation:

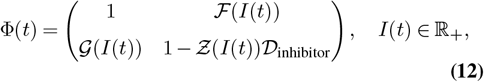

where *Ƶ* (*I*(*t*)) is given by:

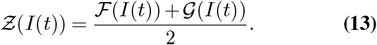

The function *Ƶ* (*I*(*t*)) captures intermediate, balanced dynamics in the presence of inhibitors by averaging transition probabilities derived from the previously-defined functions ℱ and *G*. This adjustment allows our model to account for the selective P38 inhibition of resistant-to-sensitive transitions (see SI sec. S5 for more details).

To test our model’s ability to predict phenotypic dynamics accurately under varying enza dosages and additional drug combinations, we performed model simulations calibrated against experimental conditions that differed from initial training scenarios. The experimental data obtained from a higher enza dosage regimen served as an independent test of model predictions rather than an imposed experimental constraint. The selection of varied enza dosages provided a stringent assessment of the computational model’s predictive robustness. Moreover, because experimental trajectories are single replicates and inherently noisy, dose-dependent parameter calibration further improved model reliability and predictive accuracy.

Guided by these model predictions, we experimentally validated the impact of P38 MAPK inhibition on resistant subpopulations. Co-cultures containing enza-sensitive (EGFP-labeled) and enza-resistant (mCherry-labeled) LNCaP cells were treated sequentially according to distinct drug schedules: (i) DMSO followed by enza, and (ii) DMSO followed by P38 inhibitor (SB203580). Live-cell imaging continuously monitored cellular proliferation, enabling precise quantification of differential responses. These experimental conditions (Figure 4h) served as an explicit test of our model predictions. Experimental outcomes closely matched model predictions, confirming that SB203580 substantially reduced growth specifically in enza-resistant cells while minimally impacting sensitive cells. Consequently, the resistant-to-sensitive cell ratio consistently decreased over time, validating our initial hypothesis that P38 inhibition selectively targets memory-driven resistance mechanisms.

Following these experimental validations, we further refined our model parameters specifically for combined enza and P38 inhibitor scenarios. Given the additive inhibitory effect observed experimentally, the resistant cell growth rate under combined treatment was modeled as *r*_*r,{*enza,P38*}*_ = *r*_*r*,enza_ −*r*_*r*,P38_. For the relatively brief experimental duration (approximately 12 days), sensitive cell growth rates under combined enza and P38 inhibitor treatment were effectively indistinguishable from their rates under enza alone (*r*_*s*, {enza,P38}_ = *r*_*s*,enza_ + *ϵ*). In scenarios involving P38 inhibitor administration without concurrent enza, sensitive cell growth rates defaulted to baseline DMSO control conditions (*r*_*s*,DMSO_). The minimal parameter adjustments used above represent a natural choice for enabling efficient fitting to the new P38 inhibitor data (further details are provided in the Methods).

Using the above-updated parameters and explicit incorporation of P38 inhibition into our model, we then repeated numerical simulations to predict the effects of combined enza and SB203580 administration. In particular, we modeled the dynamical effects of enza, followed by combined enza+SB203580 therapy. Our large-scale numerically predicted results (Figure 4i, blue trajectories) supported the role of enza+SB203580 co-administration in effectively preventing the outgrowth of resistant subpopulations. Lastly, follow-up experimental validation (Figure 4i, orange trajectory) provided a close alignment with our model predictions. Collectively, our results support the therapeutic potential of synergizing enza and P38 MAPK inhibition to delay resistance in PCa, in addition to the role of *in silico*-guided therapeutic scheduling.

### Validation and Clinical Application of the Computational Model

Given the findings described above, we next aimed to apply our computational model to clinical data. By integrating prostate-specific antigen (PSA) measurements from patients enrolled in a trial of intermittent androgen deprivation therapy (ADT), we sought to determine how accurately our memory-driven adaptation model could predict patientspecific tumor progression and the evolution of therapeutic resistance in clinically relevant scenarios. We applied our model to a total of 72 patients undergoing intermittent ADT and partitioned based on whether they progressed (three consecutive PSA increases above 4.0 *µ*g/L with castrate levels of serum testosterone (See Methods section for details of the clinical dataset. Intermittent treatment was suspended when PSA dropped below 4.0 *µ*g/L and resumed when PSA rose to 10*µ*g/L. This intermittent treatment policy provided a consistent framework that we could then directly implement as rule-based actions in our computational framework. We initialized the model by pre-training on a small subset (≈20%) of patients selected at random, using a uniform random sampling procedure without replacement to ensure representative coverage of both progression and non-progression cases. Then, to account for individual patient variability in tumor phenotype and its consequent impact on parameters (e.g., tumor growth rates), we calibrated the model using the earliest 10-15% of PSA measurements. This time interval was selected to capture PSA dynamics during the initial response and rebound phase and to evaluate the prognostic value of an approach intended to predict the long-term patient-specific dynamics from an initial calibration. The calibrated model was then coupled with a model-free Soft Actor–Critic (SAC) agent to learn adaptive dosing policies within this personalized environment, enabling long-term predictions of patientspecific PSA trajectories under optimized regimens.

### Computational Modeling and Simulation of Clinical PCa Dynamics

In this section, we utilized detailed longitudinal PSA measurements from patients treated with intermittent ADT to precisely calibrate and validate patient-specific computational models. Using these calibrated models, we simulated individual tumor-cell population dynamics under clinically relevant ADT conditions. An important goal of this analysis was to evaluate whether the memory-driven framework can capture both clinical patterns observed in the cohort: (i) patients with progressively shorter intermittent ADT cycles, reflecting gradual resistance emergence, and (ii) patients without shortening, who maintain more stable cycle lengths and delayed resistance. Framing the problem in this way sets up a test of whether the model can account for heterogeneous adaptive dynamics across the patient population. We explicitly captured differences between patients who eventually progressed under intermittent ADT—those showing earlier PSA rebounds and faster loss of control—and patients who remained progression-free over the follow-up period. These simulations predicted clear differences in the underlying model features. For example, progression-free cases were predicted to have a relatively small baseline fraction of androgen-resistant cells, whereas patients with progression exhibited larger pre-existing resistant subpopulations that drove earlier therapeutic failure. (SI Section S6; Figures S6–S8 for patients without progression and Figures S9–S10 for patients with progression).

Patients treated with intermittent ADT in the trial data exhibited two predominant clinical phenotypes. Progression-free cases showed a marked initial decline in PSA, likely reflecting rapid depletion of the androgen-sensitive subpopulation, followed by a delayed rebound as resistant cells gradually expanded in proportion. In contrast, patients with observed progression displayed shorter control intervals and faster PSA rebounds across successive cycles, consistent with a higher pre-existing fraction of androgen-resistant or neuroendocrine-like cells. Our calibrated model accurately reproduces these distinct clinical trajectories (Figure 5, Figures S6–S10), and simulations predict clear differences in the fractions and proliferation rates of sensitive and resistant subpopulations driving each pattern.

**Fig. 5.**
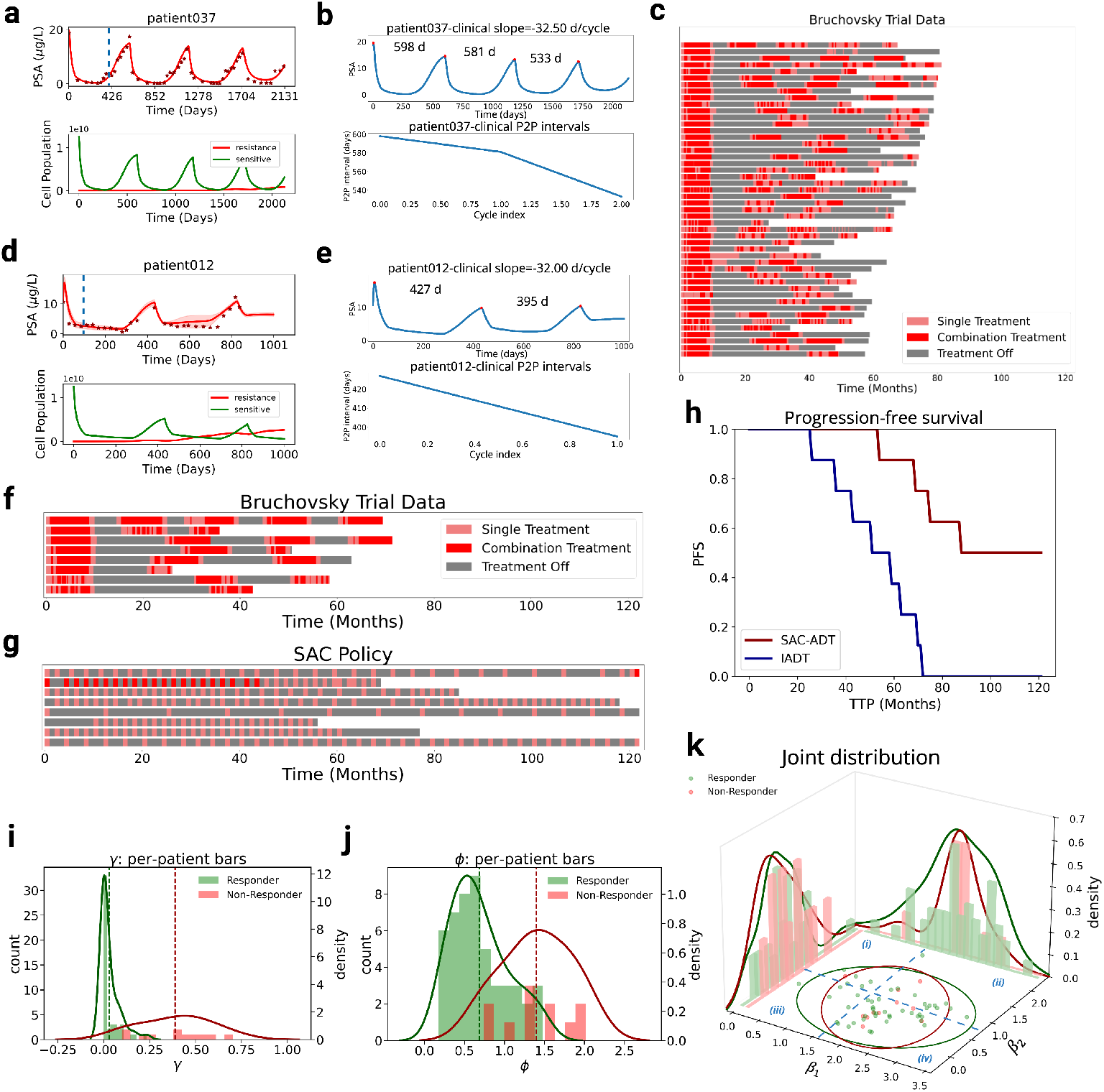
Model agreement with treatment outcomes under intermittent ADT and model-predicted adaptive policies for prostate cancer. **(a; top)** PSA trajectory for *Patient037* (without progression); predicted PSA (solid line) vs. clinically observed PSA (red dots). A vertical dashed blue line marks the calibration window (first 10–15% of the trajectory, spanning PSA decline and rebound), to the left of which the data are used for model fitting. The calibrated model is then used to predict data to the right of the vertical line. **(a; bottom)** Model-predicted population dynamics of enza-sensitive (green) and enza-resistant (red) subpopulations. **(b)** Cycle-wise contraction for *Patient037* : successive intermittent ADT response periods shorten over time, reflecting diminishing control. **(c)** Clinical intermittent ADT schedule from the Bruchovsky *et al*. trial. **(d; top)** PSA trajectory for *Patient012* (with early progression); predicted PSA (solid line) vs. clinically observed PSA (red dots) showing early loss of control. A vertical dashed blue line again denotes the calibration window, to the left of which the data are used for model fitting and to the right of which model predictive accuracy is assessed. **(d; bottom)** Model-predicted population dynamics of enza-sensitive (green) and enza-resistant (red) subpopulations. **(e)** Rapid cycle-wise contraction for *Patient012*, indicating early progression. **(f)** Clinical intermittent ADT schedule (standard). **(g)** SACADT (RL-derived) policy across progressed patients, illustrating personalized treatment intervals. **(h)** Progression-free survival (PFS) under the computationally optimized SAC-ADT prediction (red) vs. clinically observed intermittent ADT (blue) calculated for patients that underwent clinical progression; SAC-derived policies are predicted to maintain *>*50% progression-free beyond 120 months, compared to 100% progression by 60 months under intermittent ADT. **(i)** Distribution of *γ* inferred from clinical data, representing the rate at which memory drives sensitive-to-resistant transitions; patients with progression (red) show earlier shifts. **(j)** Distribution of *ϕ* inferred from clinical data, which governs PSA decay rate; progressed patients (red) exhibit faster PSA rebound. **(k)** Joint distribution of *β*_1_ and *β*_2_ inferred from clinical data, representing the strength of ADT effects on sensitive and resistant cells, respectively; patients without progression show stronger drug effects on both subpopulations, with sensitive *>* resistant (*β*_1_ *> β*_2_).

In addition to these distinct progression-related phenotypes, clinical data from the Bruchovsky *et al*. trial also revealed a consistent shortening of the off-treatment intervals across successive intermittent ADT cycles. Specifically, the mean time off therapy decreased from approximately 64 weeks in Cycle 1 to about 26 weeks by Cycle 5, a trend observed in both patients with and without progression. While those without progression maintained relatively longer off-treatment phases compared to patients who progressed, all patients exhibited progressive cycle shortening, reflecting the gradual acceleration of PSA rebound and the emergence of resistant subpopulations over time. Importantly, this progressive cycle shortening observed clinically is consistent with a memory effect, in which prior androgen suppression selects for resistant clones or activates resistant signaling pathways at the single-cell level, ultimately driving the transition to castration-resistant prostate cancer (CRPC). This effect is more gradual among patients without progression, where cycle shortening accumulates over successive treatment rounds as resistant subclones slowly expand. By contrast, patients with early progression exhibit a more abrupt loss of control, bypassing this gradual memory process. In our model, this behavior arises from higher baseline resistant fractions inferred at therapy initiation, which drive rapid PSA rebound despite treatment.

Building on this patient-specific model calibration, we next applied our model with three objectives in mind: (i) *accurate forward prediction* of each patient’s PSA course using only a short initial calibration window, (ii) *quantitative inference* of the phenotypic dynamics that underlie individual responses, and (iii) *optimization of treatment schedules*. To achieve this final goal, we embedded the calibrated tumor model in a model-free reinforcement-learning framework (Soft Actor–Critic, SAC), allowing the policy and value networks to learn adaptive dosing strategies tailored to each patient’s evolving tumor state and thereby explore personalized regimens that could extend therapeutic benefit or delay resistance.

### Memory-driven model accurately captures individual patient time course trajectories

#### RL-Guided Adaptive Therapeutic Strategies for Patients Without Progression

When applied to the cohort of progression-free patients during the observation period, the memory-driven model predicted that individuals with a high baseline fraction of enza-sensitive cells exhibited consistently low transition rates toward the resistant phenotype throughout follow-up (median ∼6 years) (Figures S6–S10). Consequently, the persistence of a large sensitive subpopulation in our model was predicted to delay the expansion of resistant cells, leading to a prolonged period of tumor control and sustained PSA suppression over extended time frames. Figure 5a illustrates this behavior for a representative progression-free case (*Patient037*; red dots denote clinical PSA values) alongside the simulated evolution of sensitive and resistant compartments. Across these patients, our model predicted that resistance appeared only after a prolonged latency—typically ∼14 months after therapy initiation—with modest inter-patient variability. While our model infers that this pattern is primarily explained by lower transition rates from sensitive to resistant states, we note that PSA dynamics alone cannot fully disentangle whether this reflects fewer phenotypic transitions or slower intrinsic growth of resistant populations. Nevertheless, the observed delayed resistance is consistent with adaptive, memory-driven phenotypic transitions rather than the rapid outgrowth of pre-existing resistance.

Figure 5b quantifies PSA cycle shortening across successive intermittent ADT cycles in a representative patient (*Patient037*) showing gradual loss of ADT control. Progressive cycle shortening is a well-documented clinical phenomenon under intermittent ADT (52), widely recognized as a hallmark of emerging castration resistance. Thus, the model’s ability to reproduce this behavior reinforces its biological relevance. Panels 5i–k summarize cohort-level parameter distributions. In particular, patients without progression exhibit higher *β*_1_ and *β*_2_ (stronger drug effects on sensitive and resistant phenotypes, respectively) and lower *γ* (slower ecological drift toward resistance) and *ϕ* (slower PSA clearance, resulting in smoother rebound dynamics), which is consistent with the prolonged control observed in Figure 5a.

Figure 5c shows the treatment schedules implemented for the intermittent ADT cohort, following the protocol of Bruchovsky *et al*. In this regimen, treatment was suspended when PSA levels fell below 4 *µ*g/L and resumed once PSA exceeded 10 *µ*g/L (52). Intermittent ADT is a threshold-based strategy in which treatment cycling indirectly fosters competition between sensitive and resistant tumor populations, thereby delaying progression. Importantly, intermittent ADT should be distinguished from adaptive therapy as defined by Gatenby *et al*. (28, 30). Whereas adaptive therapy involves algorithmic, model-driven adjustment of dosing to exploit tumor eco-evolutionary dynamics, intermittent ADT relies solely on fixed PSA thresholds; the progressive shortening of cycles observed in patients reflects tumor evolutionary dynamics under these fixed rules rather than intentional adaptive modulation.

To explore whether a model-guided strategy could outperform regimens based on fixed PSA thresholds, we used each patient’s calibrated tumor model to generate an optimal dosing schedule using our reinforcement-learning (RL) framework. Figure S11a shows the resulting RL-informed policy, which adopts an ascending pattern of dose intensity and treatment-on duration across cycles. The algorithm selectively intensifies therapy when the sensitive compartment can still suppress resistant growth and eases pressure once resistant expansion risk increases—reducing overall tumor burden while postponing progression. Our model-guided results identify alternative therapeutic policies predicted to maintain tumor control longer while minimizing total drug exposure. For patients without progression, our optimization objective was to *maximize the duration of tumor control*—that is, to lengthen the interval during which PSA remains below progression thresholds—while minimizing cumulative drug burden. In contrast, for patients who ultimately progressed, the objective shifted to *maximizing time-to-progression* (TTP) and overall survival, since these tumors harbored larger preexisting resistant fractions and were clinically confirmed to relapse earlier. In the progressed cohort, our RL-derived schedules achieved a median TTP improvement of ≈50 months over standard continuous ADT (Figure 5f–h). However, heterogeneity persisted—about half of the progressed patients still reached biochemical progression within ≈100 months, whereas the remainder maintained durable control beyond the study horizon. Retrospective inspection of baseline PSA kinetics and model-inferred resistant fractions revealed that an early, steep PSA decline under standard ADT—even if partial—was a strong predictor of greater RL-mediated TTP gains, suggesting that our framework could be used at diagnosis to identify patients most likely to benefit from adaptive dosing. Together, these findings highlight the potential prognostic value of memory-driven models of adaptation. Future prospective implementations of our framework could incorporate real-time updates from longitudinal PSA measurements, enabling continuous refinement of treatment parameters to dynamically maintain optimal therapeutic control.

#### RL-Guided Adaptive Therapeutic Strategies for Patients with Progression

RL-guided adaptive therapy has demonstrated potential for delaying time to progression (TTP) in patients exhibiting therapeutic resistance. In this study, we first calibrated a mechanistic tumor dynamics model using clinical data from patients who eventually progressed under intermittent ADT. This calibrated environment model was then used to train a model-free reinforcement learning (RL) agent to explore alternative dosing strategies. By simulating patientspecific tumor evolution under different adaptive regimens, we evaluated the effectiveness of RL-guided therapy relative to standard intermittent ADT (29).

Figure 5d shows the predicted PSA trajectory for a representative patient with early progression (*Patient012*). The SAC-ADT model accurately captures the clinical PSA dynamics during the treatment period (red dots) and projects future PSA behavior (solid red line) under continued adaptive dosing. This trajectory reflects the model’s capacity to anticipate changes in tumor phenotypic composition relative to treatment history. Figure 5e displays the corresponding *cycle-wise contraction* for *Patient012*, highlighting marked shortening of response periods across cycles that accompanies early loss of control (see SI Figures S12–S16 for complete reporting across progressed patients).

Figure 5f presents the clinical treatment schedule from the Bruchovsky *et al*. trial, which followed a rule-based intermittent ADT protocol where treatment was suspended when PSA fell below 4 *µ*g/L and resumed once PSA exceeded 10 *µ*g/L. While this fixed-threshold approach captured overall population-level dynamics, it failed to account for patientspecific tumor ecology. The cohort of patients with progression underscores these limitations: their rapid disease advancement under intermittent ADT indicates that threshold-based cycling alone was insufficient to constrain resistant subpopulations. These cases therefore highlight the need for model-guided adaptive regimens that dynamically adjust treatment based on individual tumor dynamics. To address this, we used our calibrated model to predict optimal treatment schedules and corresponding tumor trajectories, directly comparing them against outcomes under the intermittent ADT protocol. Figure 5g illustrates the resulting SAC-learned dosing strategies across the progressed cohort. These policies dynamically modulate therapy intervals in response to tumor burden and phenotypic composition, enabling *in silico* schedules that balance treatment intensity and delay resistant expansion, thereby prolonging disease control.

Upon further analysis of the eight patients categorized as clinically progressed, our model predicted substantially more heterogeneous outcomes under SAC-derived treatment policies compared with those without progression. While the majority of progressed patients were predicted to exhibit improvement in progression-free survival, our results suggested that a subset of three patients would experience limited benefit, with progression occurring before 80 months. To better understand these differences, we stratified the progressed patients into two groups based on progression times exceeding 80 months (Group A, *n* = 5) or less than 80 months (Group B, *n* = 3).

Both subgroups experienced early progression (TTP *<* 60 months) under the original Bruchovsky clinical protocol (Figure 5f), where continuous drug administration was initiated whenever PSA exceeded 10 *µ*g/L. The underlying phenotypic dynamics of our model suggest that rapid progression can be attributed to expansion of preexisting resistant clones due to delayed drug administration. Under our fitted model, simulations involving a uniform high-dose strategy predicted disproportionate depletion of drug-sensitive cells. In contrast, the SAC policy applied proactive combination pulses with intensified ON-periods interspersed with OFF or single-agent intervals. Patients in model-identified Group A were predicted to exhibit delayed progression under SAC-derived scheduling through more effective control of the resistant subpopulation. These patients were characterized by relatively low baseline resistant fractions, quasi-resistant cells with residual drug sensitivity (i.e., 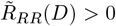), and balanced growth dynamics (e.g., comparable *K*_*G*_ and *K*_*R*_). As a result, the SAC-guided policy cyclically contracted both compartments and maintained sensitive cell dominance via the logistic crowding term. In line with adaptive therapy principles, well-timed recovery periods were predicted to allow sensitive cell regrowth, outcompeting resistant clones and delaying tumor progression well beyond clinical baselines.

Our modeling approach predicted reduced success in the Group B cohort. Closer inspection revealed that these patients possessed one or more unfavorable features: (i) a large initial burden of resistant cells (*X*_*R*_(0) ≫*X*_*G*_(0)), (ii) significantly reduced drug-mediated killing rates for resistant phenotypes, or (iii) a pronounced growth advantage of resistance (e.g., *R*_*R*_ *> R*_*G*_ and *K*_*R*_ *> K*_*G*_). In these settings, the SAC agent—though optimal in scheduling—was fundamentally constrained by the biological environment over which it was optimizing. Since the effective kill rate depends on the administered dose intensity, offset by phenotypic switching penalties (Eq. 5), patients with high switching costs and minimal baseline sensitivity were predicted to experience negligible impact even from aggressive SAC scheduling. Consequently, the dose scheduler could not meaningfully reduce the resistant compartment, and depletion of sensitive cells inadvertently accelerated progression.

Figure S11b highlights a challenging case (*Patient092*) where the clinical intermittent ADT policy failed to maintain PSA control. In contrast, the SAC-derived policy effectively suppressed PSA progression in this patient, demonstrating its ability to adaptively counter resistance that overwhelms standard protocols.

Finally, Figure 5h compares progression-free survival (PFS) between SAC-ADT-derived predicted PSA dynamics (red) and PSA dynamics obtained from the intermittent ADT clinical tiral (blue) for the cohort that underwent progression. The SAC-ADT method predicts the existence of alternative treatment policies that consistently maintained a higher fraction of progression-free patients throughout the 120-month simulation window. Notably, while all intermittent ADT–treated patients experienced clinical progression by month 60, our numerical simulations predict that the alternative SAC-ADT-derived treatment policies would achieve progression-free status in over 50% of the patient trajectories. Collectively our analysis, performed on retrospective clinical trial data, can well-explain the PSA dynamics of individual patients and also suggests improved treatment schedules to minimize the rate of progression. Panels 5i–k provide mechanistic context: patients with progression show lower *β*_1_, *β*_2_ (weaker drug effects on sensitive and resistant cells), higher *γ* (faster ecological drift toward resistant dominance), and higher *ϕ* (accelerated PSA clearance and rebound), together explaining the rapid loss of control observed in Figures 5d–e and the improved control achieved by SAC-ADT in Figures 5g–h. Importantly, in the joint parameter space (Figure 5k), these differences map onto the four characteristic regimes identified earlier. Patients in Region (iv), defined by high *β*_1_ but low *β*_2_, correspond to shortening-cycle trajectories with steep initial PSA declines followed by increasingly rapid rebounds. By contrast, patients in Region (ii), with moderate-to-high *β*_1_ and appreciable *β*_2_, exhibit more durable responses with stable or non-shortening cycles. This partitioning illustrates how the SAC framework adapts effectively across both unfavorable regimes (iv) and more stable regimes (ii). Such policies are predicted in this computational model to achieve improved long-term control over that of the clinically applied intermittent ADT policy.

Together, these findings underscore the potential clinical applicability of SAC-ADT as a flexible, patient-tailored treatment framework. By learning dosing strategies that dynamically respond to tumor evolution, SAC-ADT is able to identify treatment policies that may outperform conventional ADT policies.

### Parameter Profiles in Shortening and Persistent Intermittent ADT Cycles

Beyond distinguishing patients with and without progression, we examined whether the model could capture differences between those who exhibited progressive cycle shortening under intermittent ADT and those whose cycle durations remained comparatively stable. To this end, we performed a comparative analysis of parameter distributions across these two clinical patterns. The most notable distinctions between the two groups were observed in the parameters *γ* and *ϕ*, representing the rate of ecological drift toward resistance and the PSA clearance rate, respectively.

Patients with shortening cycles—typically those who later experienced progression—were estimated to have higher *γ* and *ϕ* values, consistent with faster ecological transitions toward resistance and sharper PSA rebound dynamics. These parameter shifts align with the clinical observation that cycle lengths progressively shorten as resistant clones emerge more rapidly following each treatment round. Examination of the joint distribution of drug-effect parameters (*β*_1_, *β*_2_) in Figure 5k further revealed that shortening-cycle patients were enriched in a region characterized by high *β*_1_ but low *β*_2_, corresponding to strong initial suppression of sensitive cells but limited efficacy against resistant populations. This imbalance explains the pattern of deep initial PSA declines followed by increasingly rapid rebounds across successive cycles.

In contrast, patients with stable cycle durations—predominantly those without progression—exhibited lower *γ* and *ϕ* values, corresponding to delayed ecological rebalancing and more gradual PSA trajectories. Parameter distributions for these patients concentrated in a regime with both moderate-to-high *β*_1_ and substantial *β*_2_ (Figure 5k), indicating that partial drug sensitivity among resistant clones constrained their expansion. This residual susceptibility of resistant cells, together with lower *γ* and *ϕ*, supported extended cycle durations and delayed resistance, preventing monotonic cycle contraction.

Taken together, these results suggest that clinically observed progressive cycle shortening emerges when treatment disproportionately targets sensitive cells while sparing resistant ones, particularly in cases prone to rapid ecological adaptation. Conversely, trajectories with preserved cycle length occur when resistant populations retain partial drug sensitivity and ecological transitions remain smoother, allowing more sustained tumor control. Importantly, these mechanistic distinctions highlight the prognostic potential of our framework, suggesting that parameter signatures distinguishing cycle shortening from stability could inform patient-specific treatment optimization in future clinical applications.

## Discussion

This study presents a comprehensive modeling framework that integrates stochastic phenotypic adaptation and RL to identify predicted sensitive and resistant cell population dynamics under androgen receptor blockade, in addition to solving for stochastic optimal treatment policies that prolong the time to therapeutic failure in the setting of PCa. By combining mechanistic modeling, clinical data calibration, and algorithmic policy optimization, we offer a patient-specific approach to designing adaptive treatment strategies aimed at overcoming therapeutic resistance.

We first analyzed a stochastic model of tumor evolution that captures memory-driven phenotypic transitions between drug-sensitive and drug-resistant states. Unlike “memoryless” models that assume Markovian switching, our framework explicitly incorporates temporal history, enabling it to reproduce lineage-traced cell dynamics observed experimentally under fluctuating enza exposure. This feature is crucial for modeling non-genetic resistance mechanisms that emerge through reversible adaptation, rather than through fixed genetic mutations.

Building on this mechanistic foundation, we evaluated the predictive capacity of our calibrated tumor model using experimental data from LNCaP cell populations exposed to various treatment regimens. The model incorporated the dynamics of enza-sensitive and enza-resistant subpopulations and was used to simulate responses under experimentally applied drug schedules. By comparing model predictions to observed cellular growth trajectories, we demonstrated that the model could accurately reproduce treatment-dependent dynamics and capture key features of resistance emergence. We applied our framework to patient data by calibrating tumor models using early PSA measurements to capture individual growth kinetics and treatment responses. The SAC agent then interacted with these personalized models to learn optimal drug schedules that adaptively modulate treatment intensity in response to tumor evolution. Unlike rule-based approaches such as intermittent ADT, the SAC-derived schedules dynamically responded to evolving tumor burden and phenotypic composition, leading to improvements in predicted progression-free outcomes under such policies, particularly among patients classified as having progression under standard protocols.

When evaluated in our model using data from patients with progression in the Bruchovsky *et al*. trial, SAC-guided therapy was predicted by simulation to achieve improved TTP and PFS relative to intermittent ADT. While all patients under intermittent ADT eventually progressed within 60 months in the clinical trial, our *in silico* simulations suggested the existence of treatment schedules that could maintain progression-free states beyond 120 months under SAC-derived policies. These findings suggest the potential benefit of personalized RL-guided scheduling as a strategy to enhance long-term disease control in patients predicted to display early progression under standard therapy, which would benefit from future follow-up first in preclinical trials.

Our results also predicted variability in the benefit of patients without progression to SAC treatment policies. Among the eight patients with progression analyzed, three were predicted to undergo progression prior to 80 months under the SAC-derived policy. Retrospective stratification revealed that patients exhibiting early progression may have harbored adverse tumor-intrinsic features, such as high baseline resistant cell burdens, negligible drug-induced killing rates in resistant clones, or growth advantages favoring resistance. In these cases, the SAC agent, despite learning optimal dosing within the constraints of the calibrated model, could not sufficiently suppress resistant populations. This limitation suggests a practical limit to what adaptive scheduling alone can achieve: RL can optimize within a patient-specific environment, but cannot overcome biological constraints hard-coded into tumor dynamics.

The effectiveness of the SAC policy thus remains constrained by patient-specific, tumor-intrinsic features that cannot be modified through scheduling alone. These findings highlight the importance of integrating biologically-targeted interventions alongside adaptive scheduling. For patients with intrinsically resistant tumor profiles, overcoming resistance likely requires additional approaches that restore drug sensitivity or reduce the initial resistant burden. One promising strategy is co-treatment with a P38 MAPK inhibitor, which has been shown to disrupt non-genetic resistance mechanisms, such as stress-induced survival signaling and phenotypic plasticity. Within our modeling framework, such interventions can increase the effective drug-induced kill rate on resistant clones, converting previously unresponsive tumors into partially sensitive ones. This biological modulation could enable SAC-derived adaptive schedules to achieve therapeutic control. Experimental evidence presented in this study supports this approach, demonstrating that P38 inhibition selectively impairs resistant subpopulations, thereby enhancing sensitivity to treatment.

Our framework relates phenotypic memory of prior drug exposures to future phenotypic transitions, and one consequence is that repeated exposures may result in different timescales of adaptation. By explicitly incorporating phenotypic memory directly into the model, we captured not only the gradual acceleration of rebound and cycle shortening seen in many patients, but also the stable cycle dynamics observed in others. This is possible because the calibrated patientspecific parameters anchor the environment to each individual’s early trajectory, while the memory component preserves the influence of prior treatment history. Together, these features enable accurate forward prediction across both shortening and non-shortening cases, underscoring the potential prognostic value of memory-driven models in distinguishing clinically divergent patient subgroups. The precise underlying molecular mechanisms that coordinate such changes remain to be precisely identified and tracked. One possible explanation is that androgen receptor inhibition by enza induces DNA damage (53, 54), in addition to epigenetic reprogramming. Repeated or prolonged enza exposure could lead to unrepaired DNA breaks that facilitate mutational processes and diversify clonal heterogeneity, whereas treatment breaks might partially spare cells from reaching this cumulative damage threshold. Clarifying the link between such molecular events and the modeled memory effect represents an important direction for future work and is the subject of active investigation.

Our findings are consistent with prior eco-evolutionary modeling studies of adaptive therapy (28–31), which showed that early treatment responses encode valuable prognostic information about the balance between drug-sensitive and drugresistant populations. Building on this foundation, our framework extends these approaches by explicitly incorporating *phenotypic memory*, enabling the model to capture history-dependent adaptation and feedback between treatment exposure and phenotypic switching. Once trained across a cohort, the model requires calibration using only the earliest 10–15% of each patient’s PSA trajectory—spanning the initial decline and rebound phases—and from this can very accurately predict long-term PSA dynamics. This striking capability demonstrates that integrating memory effects can significantly enhance both the predictive and potentially prognostic power of eco-evolutionary models of therapeutic response.

An important limitation of our modeling framework arises from its reliance on retrospective clinical cohorts. First, several complexities of the clinical trial, for example, the initial treatment with anti-androgen, lack an explicit representation in our deep learning model. The purpose of this foundational model was to establish the utility of incorporating memory-driven adaptation for reliably tracking disease dynamics over time in response to therapies. Despite its simplicity, the model performed quite well at recapitulating known trajectories using a minority of training data. Nonetheless, future efforts to add in additional pertinent clinical details are expected to enhance model applicability, provided the necessary training data exist to train these more complicated approaches. While our SAC-derived policies generated patientspecific treatment schedules and forward PSA predictions that outperformed the original clinical protocol, we were unable to implement and evaluate these strategies in real-time clinical settings. This restricted our ability to iteratively refine the model predictions based on evolving patient responses. Moreover, the absence of prospective validation remains an inherent limitation in the current work. Lastly, the SAC-derived policies were constrained to clinically feasible dosing schedules, which, while ensuring medical plausibility, may limit exploration of the broader action space considered during training. Future studies involving prospective clinical trials will be crucial to assess real-world feasibility, validate predictive accuracy, and enable continuous updating of treatment policies informed by ongoing patient data.

Taken together, these results propose a mechanistically-grounded pipeline for adaptive therapy design. The integration of memory-aware stochastic modeling, RL-based treatment optimization, and biologically-informed interventions allows for nuanced, patient-specific strategies that are predicted to extend therapeutic control. Traditional approaches based on maximally tolerated dose (MTD) regimens can inadvertently accelerate resistance by eliminating sensitive cells and providing a competitive advantage to resistant clones. By contrast, our framework leverages tumor ecoevolutionary dynamics and phenotypic memory to balance selective pressures more strategically. This contrast highlights the potential of memory-driven adaptive scheduling to address a key limitation of conventional MTD strategies. While the SAC policy proved effective in many cases, its limitations in the face of intrinsic resistance highlight the importance of combining scheduling with molecular targeting. Future extensions may incorporate real-time biomarker feedback, multi-drug environments, or Bayesian policy adaptation to further personalize treatment under uncertainty.

In conclusion, our study provides a scalable and generalizable framework for understanding the underlying dynamics of phenotypic plasticity in promoting resistance. We anticipate that our modeling frameworks will be more broadly useful for exploring improved therapeutic strategies in future treatment cohorts, in addition to understanding the role of memory-driven phenotypic adaptation in other cancer subtypes.

## Author contributions

ZSG, JAS, and JTG designed the study. JAS and JTG supervised the study. SD, PDO, and JAS developed the experimental approach. SD and PDO performed the experiments. ZSG and JTG developed the computational framework. ZSG performed the computational analysis. ZSG, SD, PDO, AJA, JAS, and JTG analyzed the results. ZSG, SD, JAS, and JTG wrote the first draft. ZSG, SD, PDO, AJA, JAS, and JTG reviewed and edited the manuscript. All authors approve of the final version of the manuscript.

## Acknowledgment

Portions of this research were conducted with the advanced computing resources provided by Texas A&M High Performance Research Computing. JTG was supported by the Cancer Prevention Research Institute of Texas (RR210080) and the National Institute of General Medical Sciences of the NIH (R35GM155458). JTG is a CPRIT Scholar in Cancer Research. JAS is supported by the National Cancer Institute (5R01CA279034).

## References

[1] Ana Bela Sarmento-Ribeiro, Andreas Scorilas, Ana Cristina Gonçalves, Thomas Efferth, and Ioannis P Trougakos. The emergence of drug resistance to targeted cancer therapies: Clinical evidence. Drug Resistance Updates, 47:100646, 2019.

[2] Aaron N Hata, Matthew J Niederst, Hannah L Archibald, Maria Gomez-Caraballo, Faria M Siddiqui, Hillary E Mulvey, Yosef E Maruvka, Fei Ji, Hyo-eun C Bhang, Viveksagar Krishnamurthy Radhakrishna, et al. Tumor cells can follow distinct evolutionary paths to become resistant to epidermal growth factor receptor inhibition. Nature medicine, 22(3):262–269, 2016.

[3] Yoh Iwasa, Martin A Nowak, and Franziska Michor. Evolution of resistance during clonal expansion. Genetics, 172(4):2557–2566, 2006.

[4] Christina Scheel, Tamer Onder, Antoine Karnoub, and Robert A Weinberg. Adaptation versus selection: the origins of metastatic behavior. Cancer research, 67(24):11476–11480, 2007.

[5] Kristel Kemper, Pauline L de Goeje, Daniel S Peeper, and Renée van Amerongen. Phenotype switching: tumor cell plasticity as a resistance mechanism and target for therapy. Cancer research, 74(21):5937–5941, 2014.

[6] Honghong Zhang, Rhonda L Brown, Yong Wei, Pu Zhao, Sali Liu, Xuan Liu, Yu Deng, Xiaohui Hu, Jing Zhang, Xin D Gao, et al. Cd44 splice isoform switching determines breast cancer stem cell state. Genes & development, 33(3-4):166–179, 2019.

[7] Paras Jain, Maalavika Pillai, Atchuta Srinivas Duddu, Jason A Somarelli, Yogesh Goyal, and Mohit Kumar Jolly. Dynamical hallmarks of cancer: Phenotypic switching in melanoma and epithelial-mesenchymal plasticity. In Seminars in Cancer Biology, volume 96, pages 48–63. Elsevier, 2023.

[8] Muhammad Sufyan Bin Masroni, Kee Wah Lee, Victor Kwan Min Lee, Siok Bian Ng, Chao Teng Law, Kok Siong Poon, Bernett Teck-Kwong Lee, Zhehao Liu, Yuen Peng Tan, Wee Ling Chng, et al. Dynamic altruistic cooperation within breast tumors. Molecular Cancer, 22(1):206, 2023.

[9] Hamzeh Kayhanian, William Cross, Suzanne EM van der Horst, Panagiotis Barmpoutis, Eszter Lakatos, Giulio Caravagna, Luis Zapata, Arne Van Hoeck, Sjors Middelkamp, Kevin Litchfield, et al. Homopolymer switches mediate adaptive mutability in mismatch repair-deficient colorectal cancer. Nature Genetics, 56(7):1420–1433, 2024.

[10] Charles C Bell and Omer Gilan. Principles and mechanisms of non-genetic resistance in cancer. British journal of cancer, 122(4):465–472, 2020.

[11] Jingxin Li, Pavithran T Ravindran, Aoife O’Farrell, Gianna T Busch, Ryan H Boe, Zijian Niu, Sean Woo, Margaret C Dunagin, Naveen Jain, Yogesh Goyal, et al. Ap-1 mediates cellular adaptation and memory formation during therapy resistance. bioRxiv, 2024.

[12] Jason T George and Herbert Levine. Stochastic modeling of tumor progression and immune evasion. Journal of theoretical biology, 458:148–155, 2018.

[13] Jason T George and Herbert Levine. Implications of tumor–immune coevolution on cancer evasion and optimized immunotherapy. Trends in Cancer, 7(4):373–383, 2021.

[14] Sydney M Shaffer, Margaret C Dunagin, Stefan R Torborg, Eduardo A Torre, Benjamin Emert, Clemens Krepler, Marilda Beqiri, Katrin Sproesser, Patricia A Brafford, Min Xiao, et al. Rare cell variability and drug-induced reprogramming as a mode of cancer drug resistance. Nature, 546(7658):431–435, 2017.

[15] Sydney M Shaffer, Benjamin L Emert, Raúl A Reyes Hueros, Christopher Cote, Guillaume Harmange, Dylan L Schaff, Ann E Sizemore, Rohit Gupte, Eduardo Torre, Abhyudai Singh, et al. Memory sequencing reveals heritable single-cell gene expression programs associated with distinct cellular behaviors. Cell, 182(4):947–959, 2020.

[16] Benjamin L Emert, Christopher J Cote, Eduardo A Torre, Ian P Dardani, Connie L Jiang, Naveen Jain, Sydney M Shaffer, and Arjun Raj. Variability within rare cell states enables multiple paths toward drug resistance. Nature biotechnology, 39(7):865–876, 2021.

[17] Kevin S Farquhar, Daniel A Charlebois, Mariola Szenk, Joseph Cohen, Dmitry Nevozhay, and Gábor Balázsi. Role of network-mediated stochasticity in mammalian drug resistance. Nature communications, 10(1):2766, 2019.

[18] Eduardo A Torre, Eri Arai, Sareh Bayatpour, Connie L Jiang, Lauren E Beck, Benjamin L Emert, Sydney M Shaffer, Ian A Mellis, Mitchell E Fane, Gretchen M Alicea, et al. Genetic screening for single-cell variability modulators driving therapy resistance. Nature Genetics, 53(1):76–85, 2021.

[19] Yogesh Goyal, Gianna T Busch, Maalavika Pillai, Jingxin Li, Ryan H Boe, Emanuelle I Grody, Manoj Chelvanambi, Ian P Dardani, Benjamin Emert, Nicholas Bodkin, et al. Diverse clonal fates emerge upon drug treatment of homogeneous cancer cells. Nature, 620(7974):651– 659, 2023.

[20] Sui Huang. Genetic and non-genetic instability in tumor progression: link between the fitness landscape and the epigenetic landscape of cancer cells. Cancer and Metastasis Reviews, 32:423–448, 2013.

[21] Sreenath V Sharma, Diana Y Lee, Bihua Li, Margaret P Quinlan, Fumiyuki Takahashi, Shyamala Maheswaran, Ultan McDermott, Nancy Azizian, Lee Zou, Michael A Fischbach, et al. A chromatin-mediated reversible drug-tolerant state in cancer cell subpopulations. Cell, 141 (1):69–80, 2010.

[22] Mohammad Fallahi-Sichani, Verena Becker, Benjamin Izar, Gregory J Baker, Jia-Ren Lin, Sarah A Boswell, Parin Shah, Asaf Rotem, Levi A Garraway, and Peter K Sorger. Adaptive resistance of melanoma cells to raf inhibition via reversible induction of a slowly dividing de-differentiated state. Molecular systems biology, 13(1):905, 2017.

[23] Jason T George and Herbert Levine. Optimal cancer evasion in a dynamic immune microenvironment generates diverse post-escape tumor antigenicity profiles. Elife, 12:e82786, 2023.

[24] Alexander Roesch, Mizuho Fukunaga-Kalabis, Elizabeth C Schmidt, Susan E Zabierowski, Patricia A Brafford, Adina Vultur, Devraj Basu, Phyllis Gimotty, Thomas Vogt, and Meenhard Herlyn. A temporarily distinct subpopulation of slow-cycling melanoma cells is required for continuous tumor growth. Cell, 141(4):583–594, 2010.

[25] Florian Rambow, Aljosja Rogiers, Oskar Marin-Bejar, Sara Aibar, Julia Femel, Michael Dewaele, Panagiotis Karras, Daniel Brown, Young Hwan Chang, Maria Debiec-Rychter, et al. Toward minimal residual disease-directed therapy in melanoma. Cell, 174(4):843–855, 2018.

[26] Piyush B Gupta, Christine M Fillmore, Guozhi Jiang, Sagi D Shapira, Kai Tao, Charlotte Kuperwasser, and Eric S Lander. Stochastic state transitions give rise to phenotypic equilibrium in populations of cancer cells. Cell, 146(4):633–644, 2011.

[27] Pujan Shrestha, Zahra S Ghoreyshi, and Jason T George. How modulation of the tumor microenvironment drives cancer immune escape dynamics. Scientific Reports, 15(1):7308, 2025.

[28] Robert A Gatenby, Ariosto S Silva, Robert J Gillies, and B Roy Frieden. Adaptive therapy. Cancer research, 69(11):4894–4903, 2009.

[29] Renee Brady-Nicholls, John D Nagy, Travis A Gerke, Tian Zhang, Andrew Z Wang, Jingsong Zhang, Robert A Gatenby, and Heiko Enderling. Prostate-specific antigen dynamics predict individual responses to intermittent androgen deprivation. Nature communications, 11(1): 1750, 2020.

[30] Jingsong Zhang, Jessica J Cunningham, Joel S Brown, and Robert A Gatenby. Integrating evolutionary dynamics into treatment of metastatic castrate-resistant prostate cancer. Nature communications, 8(1):1816, 2017.

[31] Kit Gallagher, Maximilian AR Strobl, Derek S Park, Fabian C Spoendlin, Robert A Gatenby, Philip K Maini, and Alexander RA Anderson. Mathematical model-driven deep learning enables personalized adaptive therapy. Cancer Research, 84(11):1929–1941, 2024.

[32] Neil Vasan, José Baselga and David M Hyman. A view on drug resistance in cancer. Nature, 575(7782):299–309, 2019.

[33] Xuan Wang, Haiyun Zhang, and Xiaozhuo Chen. Drug resistance and combating drug resistance in cancer. Cancer drug resistance, 2(2):141, 2019.

[34] Jason T George. Optimal phenotypic adaptation in fluctuating environments. Biophysical Journal, 122(22):4414–4424, 2023.

[35] Paras Jain, Mohit Kumar Jolly, and Jason T George. How memory and adaptation cost shape cell phenotypic dynamics in response to fluctuating environments. bioRxiv, pages 2025–05, 2025.

[36] Kathryn E Ware, Santosh Gupta, Jared Eng, Gabor Kemeny, Bhairavy J Puviindran, WenChi Foo, Lorin A Crawford, R Garland Almquist, Daniella Runyambo, Beatrice C Thomas, et al. Convergent evolution of p38/mapk activation in hormone resistant prostate cancer mediates pro-survival, immune evasive, and metastatic phenotypes. bioRxiv, pages 2020– 04, 2020.

[37] Kathryn E Ware, Beatrice C Thomas, Pelumi D Olawuni, Maya U Sheth, Nathan Hawkey, M Yeshwanth, Brian C Miller, Katherine J Vietor, Mohit Kumar Jolly, So Young Kim, et al. A synthetic lethal screen for snail-induced enzalutamide resistance identifies jak/stat signaling as a therapeutic vulnerability in prostate cancer. Frontiers in Molecular Biosciences, 10: 1104505, 2023.

[38] David Basanta, Jacob G Scott, Mayer N Fishman, Gustavo Ayala, Simon W Hayward, and Alexander RA Anderson. Investigating prostate cancer tumour–stroma interactions: clinical and biological insights from an evolutionary game. British journal of cancer, 106(1):174– 181, 2012.

[39] John T Isaacs and Donald S Coffey. Adaptation versus selection as the mechanism responsible for the relapse of prostatic cancer to androgen ablation therapy as studied in the dunning r-3327-h adenocarcinoma. Cancer research, 41(12_Part_1):5070–5074, 1981.

[40] Gouhei Tanaka, Yoshito Hirata, S Larry Goldenberg, Nicholas Bruchovsky, and Kazuyuki Aihara. Mathematical modelling of prostate cancer growth and its application to hormone therapy. Philosophical Transactions of the Royal Society A: Mathematical, Physical and Engineering Sciences, 368(1930):5029–5044, 2010.

[41] Joseph D Butner, Dalia Elganainy, Charles X Wang, Zhihui Wang, Shu-Hsia Chen, Nestor F Esnaola, Renata Pasqualini, Wadih Arap, David S Hong, James Welsh, et al. Mathematical prediction of clinical outcomes in advanced cancer patients treated with checkpoint inhibitor immunotherapy. Science advances, 6(18):eaay6298, 2020.

[42] B Ribba, Nick H Holford, Paolo Magni, I Trocóniz, I Gueorguieva, P Girard, C Sarr, M Elishmereni, C Kloft, and Lena E Friberg. A review of mixed-effects models of tumor growth and effects of anticancer drug treatment used in population analysis. CPT: pharmacometrics & systems pharmacology, 3(5):1–10, 2014.

[43] Yitao Lu, Qian Chu, Zhen Li, Mengdi Wang, Robert Gatenby, and Qingpeng Zhang. Deep reinforcement learning identifies personalized intermittent androgen deprivation therapy for prostate cancer. Briefings in Bioinformatics, 25(2):bbae071, 2024.

[44] M Ikeda and DD Šiljak. Lotka-volterra equations: Decomposition, stability, and structure: Part i: Equilibrium analysis. Journal of Mathematical Biology, 9(1):65–83, 1980.

[45] M Ikeda and DD Šiljak. Lotka-volterra equations: decomposition, stability, and structure part ii: nonequilibrium analysis. Nonlinear Analysis: Theory, Methods & Applications, 6(5): 487–501, 1982.

[46] Dalit Engelhardt. Dynamic control of stochastic evolution: a deep reinforcement learning approach to adaptively targeting emergent drug resistance. Journal of Machine Learning Research, 21(203):1–30, 2020.

[47] Sergey Ivanov and Alexander D’yakonov. Modern deep reinforcement learning algorithms. arXiv preprint 1906.10025, 2019.

[48] TP Lillicrap. Continuous control with deep reinforcement learning. arXiv preprint 1509.02971, 2015.

[49] John Schulman. Trust region policy optimization. arXiv preprint 1502.05477, 2015.

[50] Tuomas Haarnoja, Aurick Zhou, Kristian Hartikainen, George Tucker, Sehoon Ha, Jie Tan, Vikash Kumar, Henry Zhu, Abhishek Gupta, Pieter Abbeel, et al. Soft actor-critic algorithms and applications. arXiv preprint 1812.05905, 2018.

[51] John R Hershey and Peder A Olsen. Approximating the kullback leibler divergence between gaussian mixture models. In 2007 IEEE International Conference on Acoustics, Speech and Signal Processing-ICASSP’07, volume 4, pages IV–317. IEEE, 2007.

[52] Nicholas Bruchovsky, Laurence Klotz, Juanita Crook, Shawn Malone, Charles Ludgate, W James Morris, Martin E Gleave, and S Larry Goldenberg. Final results of the canadian prospective phase ii trial of intermittent androgen suppression for men in biochemical recurrence after radiotherapy for locally advanced prostate cancer: clinical parameters. Cancer, 107(2):389–395, 2006.

[53] Jonathan F Goodwin, Matthew J Schiewer, Jeffry L Dean, Randy S Schrecengost, Renée de Leeuw, Sumin Han, Teng Ma, Robert B Den, Adam P Dicker, Felix Y Feng, et al. A hormone–dna repair circuit governs the response to genotoxic insult. Cancer discovery, 3 (11):1254–1271, 2013.

[54] William R Polkinghorn, Joel S Parker, Man X Lee, Elizabeth M Kass, Daniel E Spratt, Phillip J Iaquinta, Vivek K Arora, Wei-Feng Yen, Ling Cai, Deyou Zheng, et al. Androgen receptor signaling regulates dna repair in prostate cancers. Cancer discovery, 3(11): 1245–1253, 2013.

