## Supplementary Information for "A memory-driven reinforcement learning model of phenotypic adaptation for anticipating therapeutic resistance in prostate cancer"

### S1 Detailed Explanation of Phenotypic Subpopulations and Transitions

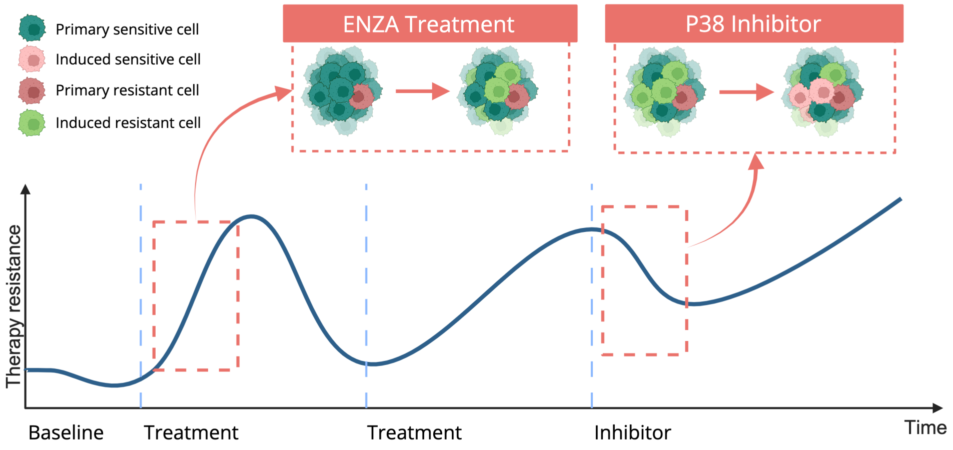


Figure S1: Schematic representation of the subpopulation dynamics and transitions under sequential therapeutic interventions. Initially, predominantly sensitive cell populations (primary sensitive) acquire resistance under selective pressures from ENZA treatment, giving rise to induced resistant cells without changing their fluorescent labeling (green cells transitioning to resistant phenotypes, $x_{RG}$). Upon subsequent exposure to the P38 inhibitor, a subset of these resistant cells can transition back to sensitivity, generating induced sensitive cells retaining their original resistance-associated labeling (red cells transitioning back to sensitivity, $x_{SR}$). These transitions highlight the phenotypic plasticity and memory-driven adaptation observed experimentally.

The computational framework presented in the main text explicitly models prostate cancer cell populations with four distinct subpopulations to capture observed experimental dynamics. These four subpopulations represent sensitive and resistant phenotypes tracked by two distinct fluorescent markers (EGFP for sensitive and mCherry for resistant). Here, we define each subpopulation and explain their biological relevance in detail:

- **Sensitive Green Cells ($x_{SG}$):** These cells represent the androgen-sensitive phenotype, initially labeled with green fluorescence (EGFP). Under drug-free or androgen-rich conditions, they proliferate rapidly due to sensitivity to androgen signaling. However, their growth significantly diminishes upon exposure to androgen deprivation therapies (ADT).
- **Resistant Green Cells ($x_{RG}$):** Cells in this category originated as sensitive green cells but acquired resistance through genetic or epigenetic modifications without losing their green fluorescent labeling. This intermediate phenotype captures the biologically critical scenario in which cells transition toward resistance but have not yet fully committed to expressing resistance-associated markers.
- **Resistant Red Cells ($x_{RR}$):** Represented by cells inherently resistant to androgen deprivation and labeled red (mCherry). These cells exhibit minimal proliferation changes upon ADT administration and can persist and expand due to their intrinsic resistance mechanisms, posing significant clinical challenges.
- **Sensitive Red Cells ($x_{SR}$):** These cells initially arose from resistant red cells but underwent transitions back to androgen sensitivity, maintaining their red fluorescent marker. This subpopulation captures the biologically relevant but less common scenario of cells reverting to sensitivity, reflecting epigenetic plasticity within tumor populations.

Experimental observations indicate that cellular transitions occur frequently between these subpopulations, yet cells retain their original fluorescent labels, suggesting phenotypic plasticity involving intermediate or transitional states. The retention of the original fluorescent marker throughout transitions emphasizes the non-instantaneous, gradual adaptive changes within each cell lineage, highlighting the dynamic complexity inherent in tumor cell evolution. Figure S1 visually illustrates these transitions, specifically demonstrating how initial *primary sensitive cells* can transition to *induced resistant cells* under ENZA treatment, and how *primary resistant cells* can revert to *induced sensitive cells* upon subsequent exposure to the P38 inhibitor.

These biologically distinct subpopulations enable our model to explicitly account for competitive dynamics and adaptive pressures observed during treatment. By modeling each subpopulation explicitly, the framework provides a refined understanding of how resistant populations gradually emerge and dominate under selective pressures imposed by androgen-targeted treatments.

The growth dynamics and transition probabilities among these subpopulations are modulated by environmental factors (e.g., drug exposure, hormonal levels) and historical cellular experiences (memory), ensuring biologically realistic simulations. This explicit modeling strategy is crucial for accurately capturing therapeutic responses observed in high temporal resolution experimental data, thus improving our ability to predict treatment outcomes and resistance development at the subpopulation level.

### S2 Detailed Analysis of Phenotype Transition Dynamics

The transitions between prostate cancer cell phenotypes under varying therapeutic conditions are governed by two memory-driven logistic functions, $\mathcal{F}(I(t))$ and $\mathcal{G}(I(t))$. These functions quantify the probabilities of transitioning from resistant to sensitive states and from sensitive to resistant states, respectively, in response to cumulative therapeutic pressures encapsulated in the memory window $I(t)$. The steepness and asymmetry of these logistic transitions are controlled by the learnable parameters $\gamma_1$ and $\gamma_2$. By adjusting these parameters during model calibration, we can accurately capture empirical observations and biologically relevant nonlinear dynamics in phenotypic adaptation.

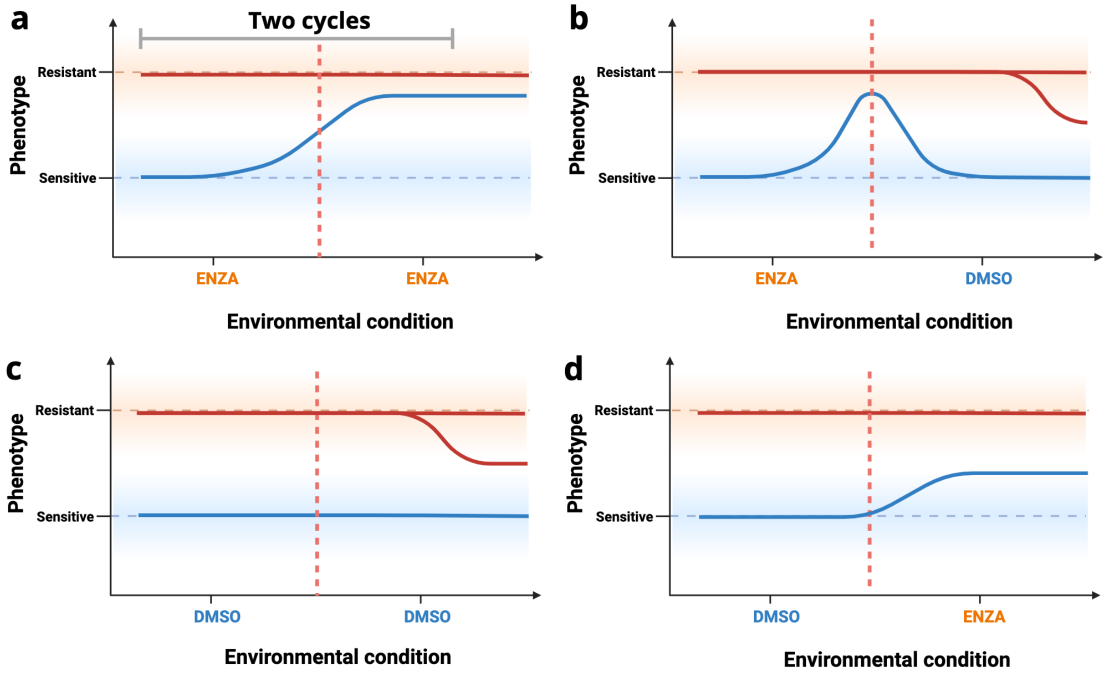


Figure S2: **Phenotypic Transition Plasticity.** (a) Under continuous ENZA treatment, sensitive cells show progressively increasing transitions toward resistance, while resistant cells remain largely fixed due to persistent suppression of $\mathcal{F}(I(t))$. (b) In the ENZA to DMSO sequence, transitions to resistance initially rise but stabilize after withdrawal, while $\mathcal{F}(I(t))$ remains low, reflecting residual effects of prior ENZA exposure. (c) Under continuous DMSO treatment, transitions are minimal for sensitive cells, while resistant cells exhibit a delayed increase in re-sensitization potential after accumulating sufficient environmental memory. (d) When DMSO precedes ENZA, sensitive cells rapidly transition to resistance upon ENZA initiation, but resistant cells, still influenced by earlier ENZA exposure, show no significant increase in $\mathcal{F}(I(t))$. These scenarios collectively demonstrate the asymmetric and memory-dependent nature of phenotypic plasticity modeled by the logistic functions.

We further illustrate these transitions across four experimentally relevant therapeutic scenarios (see Figure S2). Under continuous Enzalutamide (ENZA) treatment (Figure S2a), the probability of transition from sensitive to resistant phenotypes, captured by $\mathcal{G}(I(t))$, progressively increases as treatment pressure accumulates. In contrast, the transition probability from resistant to sensitive phenotypes, represented by $\mathcal{F}(I(t))$, remains very low from the beginning ($\mathcal{F}(I(0)) \ll 1$) and continues to decline over time. This indicates that, within an ENZA-only environment, resistant cells exhibit minimal phenotypic plasticity and are unlikely to revert to a sensitive state. This asymmetric transition pattern mirrors experimental findings, wherein sustained ENZA exposure imposes strong selective pressure that promotes the emergence and dominance of resistant phenotypes.

When ENZA administration is followed by vehicle control (ENZA, DMSO; Figure S2b), sensitive cells initially exhibit increased transitions toward resistance during ENZA exposure. Upon switching to DMSO, the absence of therapeutic pressure leads to a gradual reduction in the transition probability $\mathcal{G}(I(t))$. Meanwhile, the transition from resistant to sensitive states ($\mathcal{F}(I(t))$) stabilizes at a low level but does not recover significantly, indicating a lingering suppressive memory effect from prior ENZA exposure.

Under continuous DMSO treatment (DMSO, DMSO; Figure S2c), both phenotypes exhibit stable transition dynamics in the absence of pharmacological pressure. Sensitive cells primarily proliferate and show minimal transition to resistant states, as the DMSO environment favors their growth and does not induce phenotypic switching. Resistant cells, on the other hand, initially exhibit negligible transitions; however, after two cycle of DMSO exposure, they begin to accumulate environmental memory. This leads to a gradual increase in the probability of transitioning back to the sensitive phenotype, particularly after two consecutive cycles. These dynamics reveal a delayed but memory-driven plastic response of resistant cells in a drug-free environment, reflecting a potential for re-sensitization in the absence of selective pressure.

In the final scenario, where DMSO is followed by ENZA (DMSO, ENZA; Figure S2d), transition probabilities shift sharply in response to ENZA initiation. Sensitive cells, previously unaffected under DMSO, now show a rapid increase in $\mathcal{G}(I(t))$, indicating enhanced transitions to resistance driven by rising treatment pressure. In contrast, the transition from resistant to sensitive states ($\mathcal{F}(I(t))$) remains suppressed throughout this scenario. This lack of re-sensitization is attributed to two factors: (i) only one preceding DMSO cycle, which provides insufficient time to induce memory-driven reversion in resistant cells, and (ii) the initial state of resistant cells already shaped by prior ENZA exposure before the experimental timeline began. As a result, resistant phenotypes retain a strong memory of drug pressure, showing little to no transition back to sensitivity even after temporary withdrawal of therapy. This transition profile highlights the immediate and lasting impact of ENZA in promoting resistance and suppressing phenotypic plasticity.

Together, these analyses demonstrate that the logistic transition functions $\mathcal{F}(I(t))$ and $\mathcal{G}(I(t))$, parameterized by $\gamma_1$ and $\gamma_2$, effectively capture the evolving plasticity of prostate cancer cell phenotypes under a range of treatment regimens. Under sustained ENZA administration, transitions are heavily biased toward resistance with suppressed re-sensitization. When ENZA is followed by DMSO, transitions to resistance initially increase but gradually stabilize, while transitions from resistance to sensitivity remain low due to lingering memory effects. In the DMSO-only condition, minimal transition activity is observed, particularly for sensitive cells, whereas resistant cells may regain plasticity after prolonged treatment withdrawal. Lastly, when DMSO is followed by ENZA, we observe a sharp increase in resistance acquisition without meaningful re-sensitization. These distinct response patterns across therapeutic sequences underscore the model's ability to encode asymmetric and memory-dependent phenotype transitions. This framework provides a biologically grounded and temporally responsive tool for predicting phenotypic dynamics, offering insights that are critical for the development of adaptive and effective treatment strategies in prostate cancer.

### S3 Learning an Adaptive Dosing Policy

#### S3.1 Soft Actor-Critic (SAC) Algorithm Details

Reinforcement learning (RL) involves iterative interactions between an agent and its environment, where at each discrete time step $t$, the agent observes the state $s_t \in S$ and selects an action $a_t \in A$. Subsequently, the environment transitions to a new state $s_{t+1}$ and provides a reward $r_{t+1}$. The policy $\pi_\phi$, parameterized by $\phi$, maps states to probability distributions over actions, allowing the agent to optimize expected cumulative rewards.

The Soft Actor-Critic (SAC) algorithm enhances traditional policy gradient methods by incorporating an entropy regularization term, promoting stochastic exploration. Its objective is to maximize a modified expected return that includes an entropy term:

$$J(\pi_\phi) = \mathbb{E}_{\tau \sim \pi_\phi} \left[ \sum_{t=0}^{T} \gamma^t \left( r(s_t, a_t) + \alpha \mathcal{H}(\pi_\phi(\cdot|s_t)) \right) \right], \tag{S1}$$

where $\gamma$ is the discount factor, $\alpha$ is the temperature parameter that balances exploration and exploitation, and $\mathcal{H}(\pi_\phi(\cdot|s_t))$ is the entropy of the policy at state $s_t$. To mitigate overestimation bias in value function updates, SAC maintains two separate critic networks $Q_{\theta_1}$ and $Q_{\theta_2}$. Each critic minimizes the mean-squared error between the predicted Q-value and a soft target value:

$$J_Q(\theta_i) = \mathbb{E}_{(s_t, a_t, r_t, s_{t+1}) \sim \mathcal{D}} \left[ \left( Q_{\theta_i}(s_t, a_t) - y_t \right)^2 \right], \tag{S2}$$

where the target $y_t$ is defined as:

$$y_t = r_t + \gamma \mathbb{E}_{a_{t+1} \sim \pi_\phi} \left[ \min_{j=1,2} Q_{\hat{\theta}_j}(s_{t+1}, a_{t+1}) - \alpha \log \pi_\phi(a_{t+1} \mid s_{t+1}) \right]. \tag{S3}$$

The policy is trained to maximize the entropy-augmented Q-value, and its loss is given by:

$$J_\pi(\phi) = \mathbb{E}_{s_t \sim \mathcal{D}} \left[ \mathbb{E}_{a_t \sim \pi_\phi} \left[ \alpha \log \pi_\phi(a_t|s_t) - Q_\theta(s_t, a_t) \right] \right]. \tag{S4}$$

To automatically tune the entropy coefficient $\alpha$, SAC minimizes the following loss:

$$J(\alpha) = \mathbb{E}_{a_t \sim \pi_\phi} \left[ -\alpha \log \pi_\phi(a_t|s_t) - \alpha \mathcal{H}_{\text{target}} \right], \tag{S5}$$

where $\mathcal{H}_{\text{target}}$ is a predefined target entropy. This adaptive temperature mechanism ensures that the exploration remains sufficient throughout training.

#### S3.2 Details about States, Action Spaces, and Reward Assignment

In our system, two primary and two intermediate phenotypes of prostate cancer cells are considered. At each continuous time step $t$, the observed cell populations $x_{SG,t}$, $x_{RG,t}$, $x_{RR,t}$, and $x_{SR,t}$ represent the state vector $s_t = (x_{SG,t}, x_{RG,t}, x_{RR,t}, x_{SR,t})$. Additional informative features, including instantaneous growth and decay rates $\dot{s_t}$, are derived directly from these cell counts and reflect the underlying drug effects and phenotype transitions within the prostate cancer environment. The explicit inclusion of the time variable $t$ further enriches the state representation, leading to a comprehensive continuous state space $S_t = (s_t, \dot{s_t}, t)$.

The action space is defined continuously (Figure S3), consisting of administered doses of Enzalutamide (ENZA) and the P38 inhibitor at each time step $t$, described by the vector $A_t = (\mathcal{D}_{t,Enza}, \mathcal{D}_{t,P38})$. Following the determination of an optimal dosing strategy through the continuous RL policy, doses are discretized into clinically relevant intervals to facilitate practical administration. A successful treatment regimen is characterized by its ability to suppress resistant cancer cell populations while preserving sensitive cancer cell populations, thereby maximizing patient survival and quality of life. The optimality criterion balances prolonged survival with minimal cumulative drug exposure, ensuring effectiveness with reduced side effects.

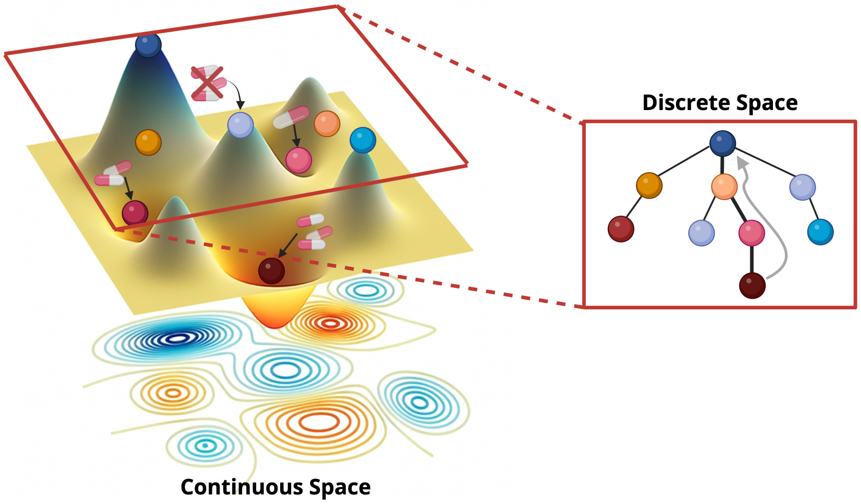


Figure S3: Schematic illustration of continuous and discrete state-action spaces in a SAC framework where the agent represents the cancer cell. In the continuous space (left), the agent navigates a smooth landscape defined by molecular and microenvironmental states, with actions representing continuous adaptation strategies in response to external signals (e.g., therapy). The lower contour plot captures the fine-grained structure of this state space. The right panel shows a discretized abstraction of this system, where both states and actions are mapped into a finite decision tree, enabling interpretable modeling and policy execution. This reflects how learned continuous strategies can be translated into discrete clinical or analytical interventions.

Leveraging the complete feature set $S_t$ in a structured, interpretable model enhances training efficiency and reward assignment accuracy. The explicit reward function incorporates multiple factors crucial to therapeutic success, including direct measures of drug efficacy and phenotype transitions. Specifically, the effectiveness of ENZA in controlling sensitive cells and the inhibitor's efficacy against resistant cells are quantified by rewards $r_{drug,t} = w_1(1 - x_{SG,t})$ and $r_{inhibitor,t} = w_2(1 - x_{RR,t})$, respectively, where $w_1$ and $w_2$ are weighting constants. Moreover, long-term disease control necessitates limiting phenotype transitions; thus, the transition reward $r_{trans,t} = w_3(1 - x_{RG,t} - x_{SR,t})$, where $w_3$ is a weighting constant, accounts for these dynamics explicitly.

To encourage intermittent dosing and avoid prolonged continuous exposure, a time-dependent penalty $p_{drug,t}$ is introduced for sustained drug administration. This penalty is structured as follows:

$$p_{drug,t} = \begin{cases} w_\alpha & \text{if } t - t_{\text{last dose}} < \delta(t), \\ 0 & \text{otherwise,} \end{cases} \tag{S6}$$

where $\delta(t) = \delta_0 e^{-\lambda t}$ represents a decaying function of time since the last dose, $\lambda$ is the decay rate, and $w_\alpha$ is a drug-specific penalty coefficient.

The cumulative step reward $r_{step,t}$, combining efficacy and penalty terms, is defined as:

$$r_{step,t} = r_{drug,t} + r_{inhibitor,t} + r_{trans,t} - p_{drug,t}. \tag{S7}$$

However, under conditions of insufficient dosing, sensitive cancer cells may proliferate rapidly and reach carrying capacity, thereby maintaining an undesirable, high-tumor state. In this scenario, the reward function alone might inadequately penalize low-dose strategies, potentially leading the RL agent toward a suboptimal, zero-dose equilibrium. To mitigate this convergence issue, we augment the reward structure by incorporating a progression-free survival bonus and integrating a metastasis probability model as an end-of-simulation (EOS) criterion. This ensures that the learned dosing strategy remains clinically relevant, effectively balances therapeutic efficacy with side effects, and avoids biologically unrealistic outcomes.

### S4 Evaluation of the Null Model Without Memory or Phenotypic Transitions

To evaluate the necessity of memory-driven dynamics and phenotypic plasticity, we constructed a null model that excludes both historical drug exposure and phenotype transitions. In this framework, each subpopulation is governed purely by intrinsic birth and death processes, with no conversion between sensitive and resistant states. The model was fit to experimental data under the ENZA–ENZA condition and then tested on ENZA–DMSO, DMSO–DMSO, DMSO, ENZA scenarios (Figure S4a–d). While the null model could partially reproduce early trends, it failed to capture key features of the adaptive response, including shifts in the EnzaR/EnzaS ratio and nonlinear changes in population composition over time. These shortcomings highlight that birth-death dynamics alone are insufficient to explain the observed behavior and confirm that incorporating both memory effects and phenotype transitions is critical for accurate modeling of drug response in heterogeneous cell populations.

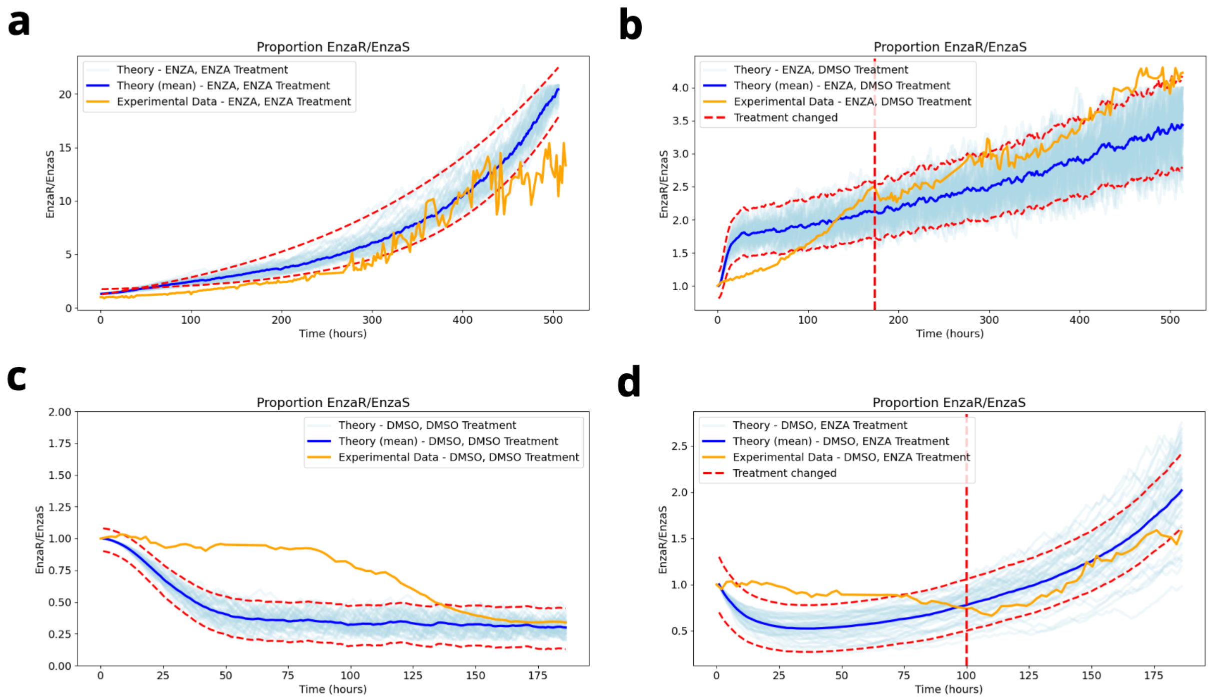


Figure S4: Simulated growth dynamics of ENZA-sensitive (green) and ENZA-resistant (red) prostate cancer cell populations predicted by the null model under various treatment regimens (orange). Panel (a) shows predicted dynamics (blue) for two consecutive cycles of ENZA treatment; panel (b) shows the ENZA–DMSO sequence; panel (c) corresponds to two cycles of DMSO treatment; panel (d) represents DMSO followed by ENZA. Unlike the memory-based model, the null model assumes fixed phenotypes and models growth solely through birth and death rates, without incorporating transitions or adaptive memory, resulting in limited capacity to replicate observed treatment-dependent adaptation.

### S5 Detailed Analysis of Phenotype Transition Dynamic - P38

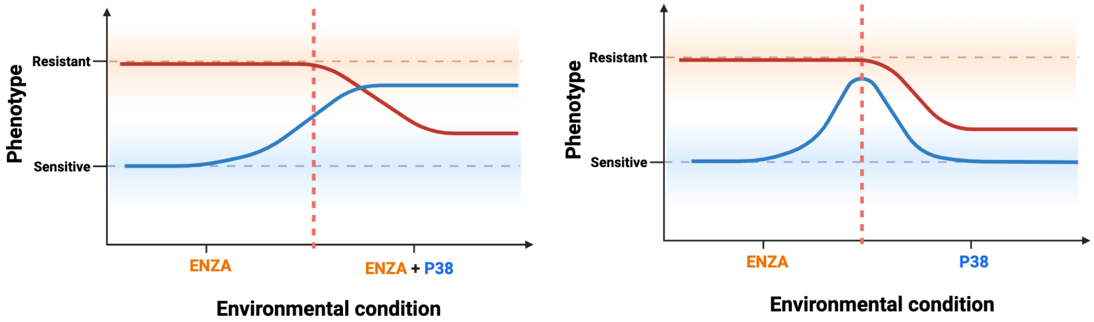


Figure S5: Schematic representation of phenotype dynamics under ENZA and P38 inhibitor treatments. (Left) When ENZA is followed by the co-administration of ENZA and P38 inhibitor, the transition from sensitive to resistant phenotype is reduced and partially reversed due to inhibitor-mediated modulation. (Right) When ENZA is followed by P38 inhibitor alone, the phenotypic shift is transient, and the resistant state partially stabilizes. These dynamics are modeled by modifying the phenotype transition matrix to include a binary inhibitor indicator and an averaged transition function $\mathcal{Z}(I(t))$, capturing the inhibitor's selective effects.

To account for the selective modulation introduced by the P38 inhibitor, we extended the phenotype transition matrix to explicitly depend on the presence or absence of the inhibitor treatment. A binary indicator variable $\mathcal{D}_{\text{inhibitor}} \in \{0, 1\}$ was defined to reflect the administration status of the P38 inhibitor. This variable was incorporated into the transition matrix $\Phi(t)$, enabling the model to downregulate transitions from resistant to sensitive phenotypes when the inhibitor is active.

The adjusted matrix includes the newly introduced function $\mathcal{Z}(I(t))$, which represents an averaged transition tendency based on the combined influence of both forward and backward transition functions $\mathcal{F}(I(t))$ and $\mathcal{G}(I(t))$. By averaging these contributions, $\mathcal{Z}(I(t))$ captures intermediate modulation of phenotype dynamics under the presence of both ENZA and P38 inhibitor.

Figure S5 illustrates this selective inhibition mechanism: when the P38 inhibitor is co-administered with ENZA (left), the transition from sensitive to resistant is reduced and partially reversed; in contrast, when the inhibitor follows ENZA (right), the phenotypic response exhibits a transient shift but ultimately stabilizes under the inhibitor-only condition. This behavior is captured through the modified transition matrix and enables the model to reflect experimentally observed resistance dynamics more accurately.

#### S6 Clinical data and PSA Level Prediction Model

To accurately represent PSA dynamics under treatment conditions, we developed a mathematical model describing serum PSA levels using a novel first-order ordinary differential equation (ODE) formulation. Specifically, our PSA dynamics are governed by the equation:

$$\frac{dP(t)}{dt} = \alpha \left[\beta_s S(t) + \beta_r R(t)\right] - \phi P(t), \tag{S8}$$

where $P(t)$ denotes the serum PSA concentration at time $t$, measured in $\mu g/L$. The terms $S(t)$ and $R(t)$ represent the androgen-sensitive and androgen-resistant tumor cell populations, respectively. Here, $\alpha$ is a scaling factor characterizing overall PSA secretion efficiency into the bloodstream, while $\beta_s$ and $\beta_r$ represent the specific PSA secretion rates from sensitive and resistant tumor cells, respectively. The parameter $\phi$ signifies the natural metabolic clearance rate of PSA from the serum, measured in $day^{-1}$.

Parameters $\alpha$, $\beta_s$, $\beta_r$, and $\delta$ were calibrated by fitting early longitudinal clinical PSA data from patients undergoing androgen-targeted therapies. Once estimated, these parameters remained fixed to predict patient-specific PSA trajectories for subsequent treatment intervals, enabling robust and accurate forecasting of tumor dynamics and therapeutic responses. This refined modeling approach effectively captures both the secretion and clearance kinetics of PSA, thereby offering improved clinical realism and predictive accuracy in evaluating tumor responses and treatment strategies.

##### S6.1 Extended Clinical PSA Predictions for Patients With and Without Progression

This section provides comprehensive model predictions of PSA trajectories for all patients under the clinical adaptive dosing policy from the Bruchovsky trial. Figures S6–S8 display predicted versus observed PSA dynamics for patients without progression, highlighting the accuracy and reliability of our computational model in capturing therapeutic responses to intermittent androgen deprivation therapy. Figures S9–S10 similarly illustrate model predictions for patients with progression, demonstrating how the clinical dosing schedules influenced PSA rebound in cases exhibiting emerging or intrinsic resistance. These predictions further validate our computational framework and reinforce its utility for simulating patient-specific responses and informing personalized treatment strategies.

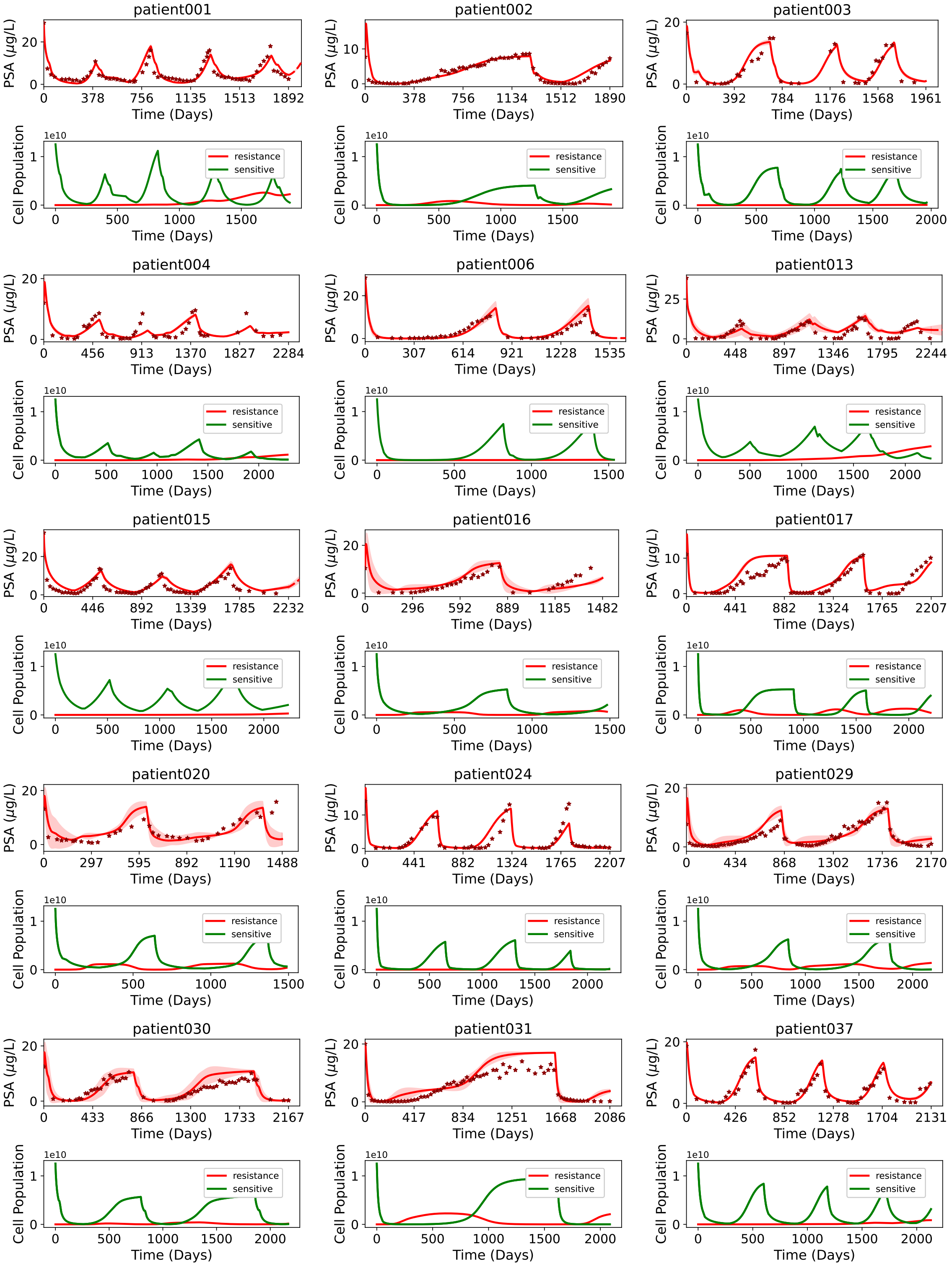


Figure S6: Model-predicted PSA trajectories (solid red lines) compared with clinical PSA measurements (red dots) for prostate cancer patients without progression (Part 1).

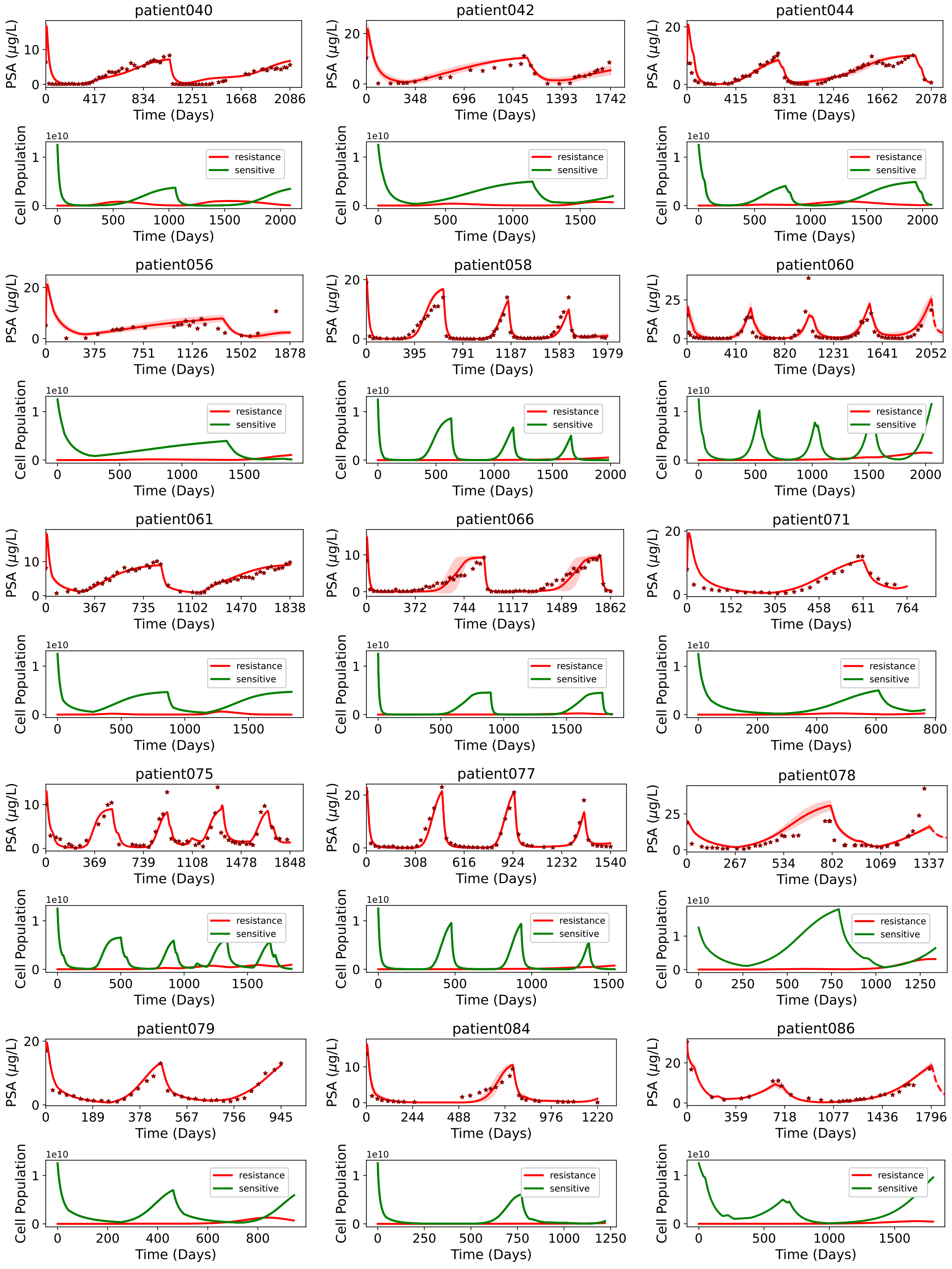


Figure S7: Model-predicted PSA trajectories (solid red lines) compared with clinical PSA measurements (red dots) for prostate cancer patients without progression (Part 2).

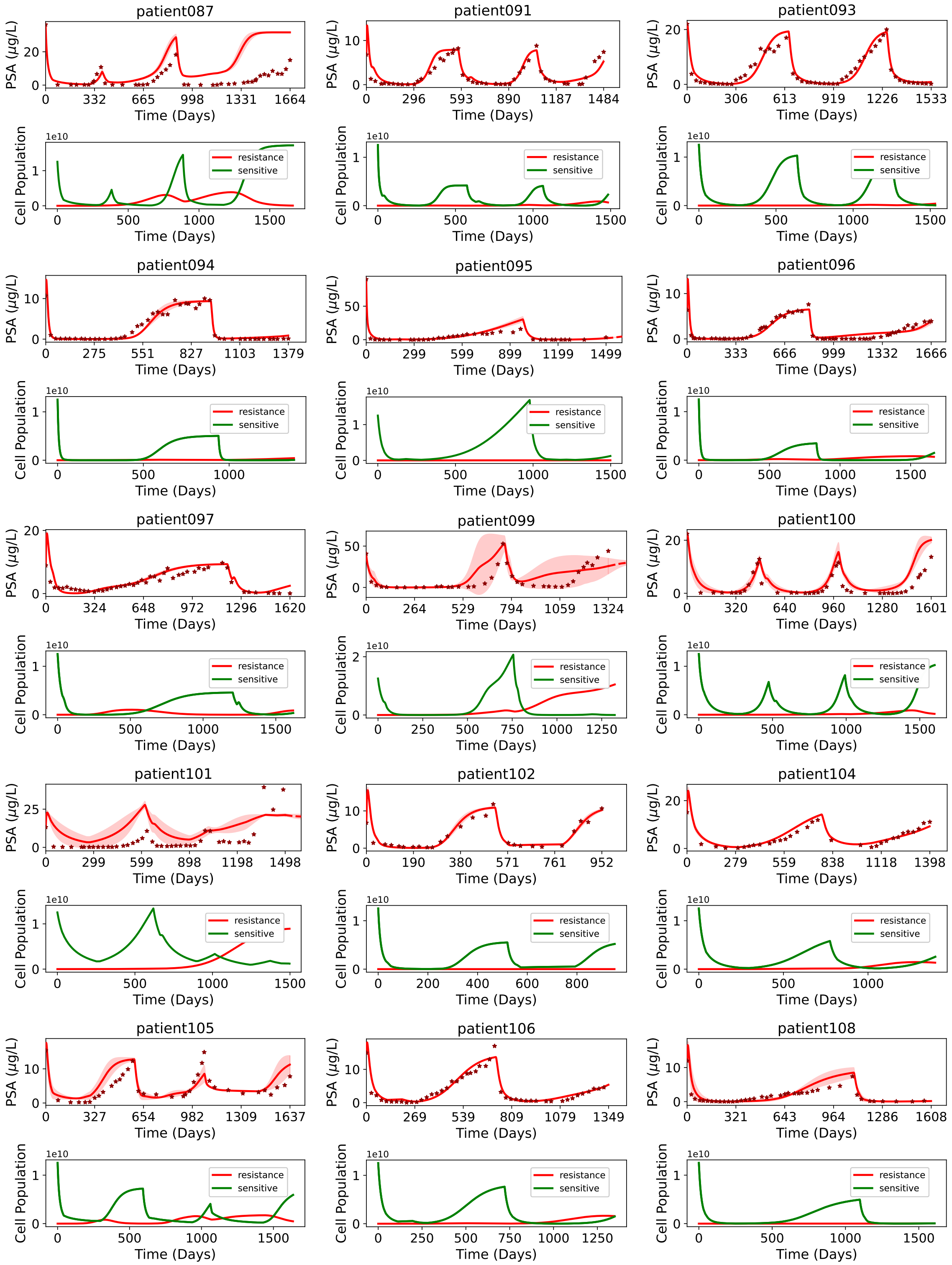


Figure S8: Model-predicted PSA trajectories (solid red lines) compared with clinical PSA measurements (red dots) for prostate cancer patients without progression (Part 3).

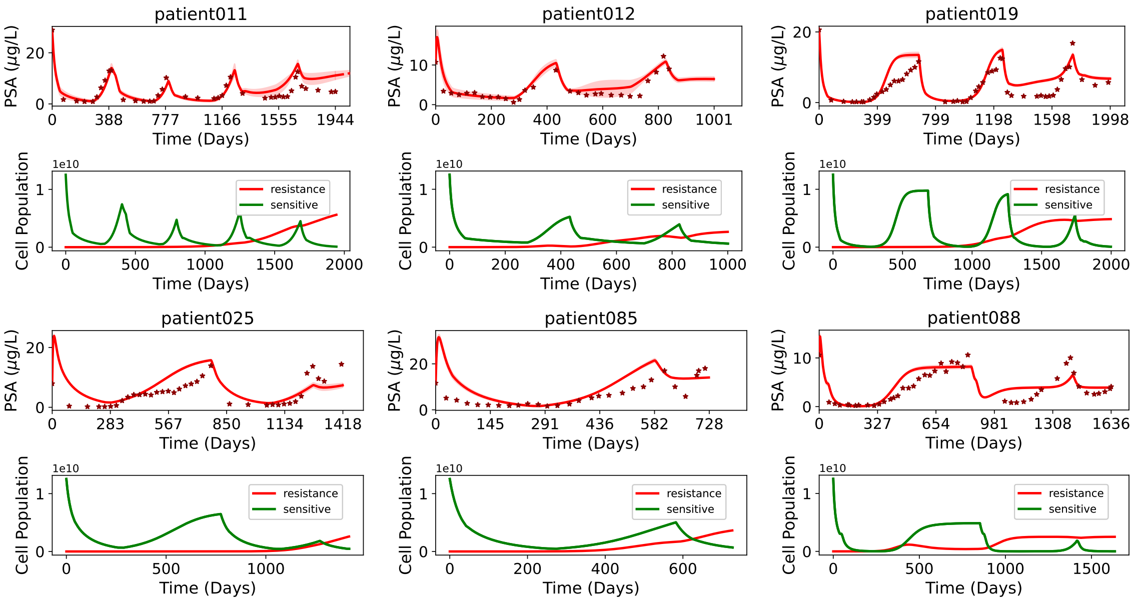


Figure S9: Model-predicted PSA trajectories (solid red lines) compared with clinical PSA measurements (red dots) for prostate cancer patients with progression.

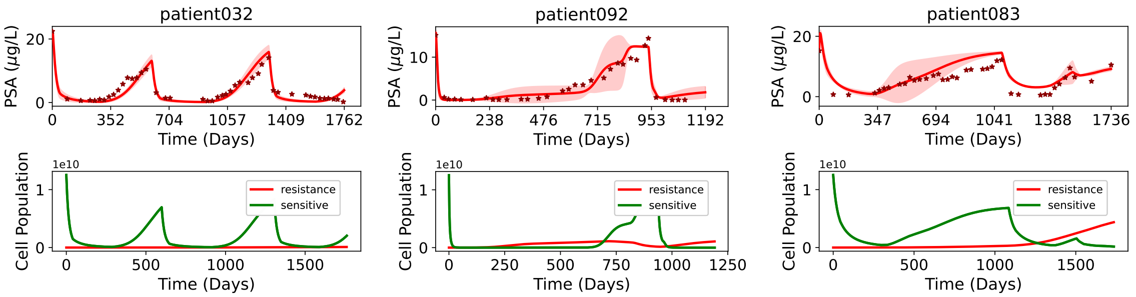


Figure S10: Model-predicted PSA trajectories (solid red lines) compared with clinical PSA measurements (red dots) for prostate cancer patients who developed metastasis.

#### S6.2 Model-Guided Adaptive Therapy Strategies Across Patients With and Without Progression

To evaluate the ability of reinforcement learning (RL) to enhance adaptive therapy, we applied our model-guided framework across both patient groups: those without progression and those with progression. Figure S11a illustrates RL-informed dosing schedules for patients without progression, which progressively adjust dose intensity and treatment-on duration to sustain tumor control while minimizing drug exposure. In contrast, Figure S11b highlights a representative case of progression (Patient092) where the clinical IADT policy failed to maintain PSA control. Here, the SAC-derived policy adaptively suppressed PSA progression, demonstrating the potential of RL-guided therapy to manage progressive disease more effectively than standard regimens.

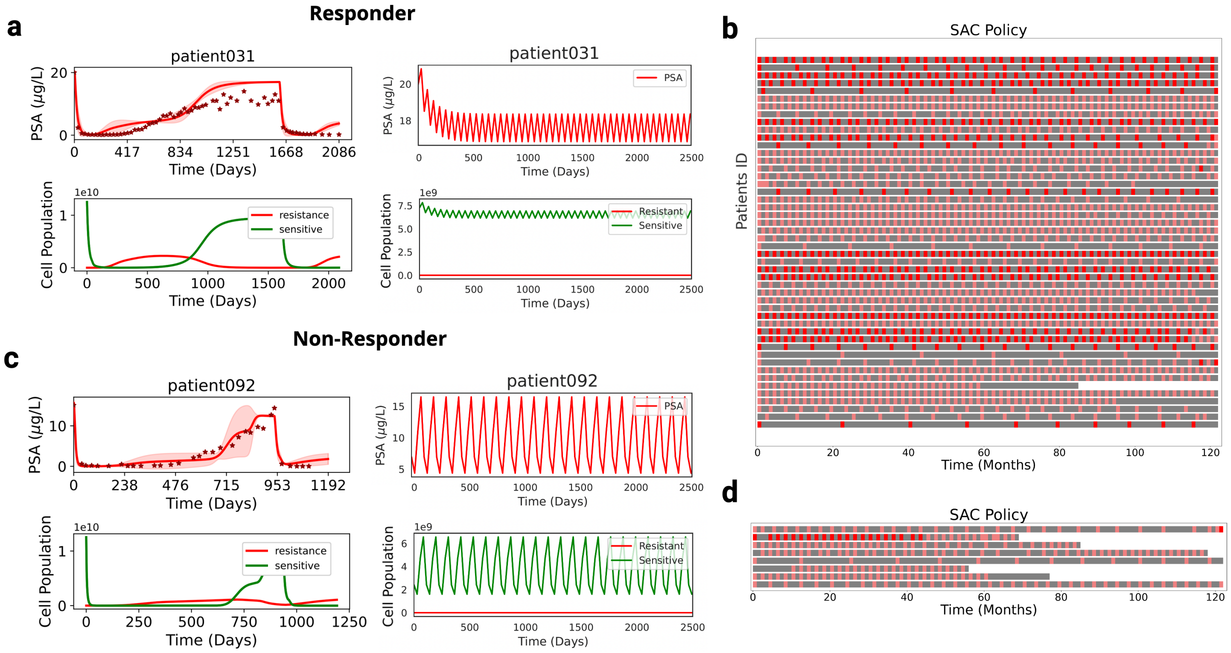


Figure S11: Reinforcement learning–guided adaptive therapy across prostate cancer patients with and without progression. (a) RL-informed dosing schedules for patients without progression, showing an ascending pattern of dose intensity and treatment-on duration across cycles. The strategy intensifies therapy when the sensitive compartment retains control over resistant clones and relaxes pressure once resistance risk increases, thereby maintaining tumor control with lower overall drug exposure. (b) PSA trajectory for a patient with progression (Patient092). While the clinical IADT policy failed to suppress PSA progression, the SAC-derived policy adaptively maintained long-term PSA control, underscoring its potential to delay resistance where standard approaches falter.

### S7 Peak-to-Peak PSA Levels

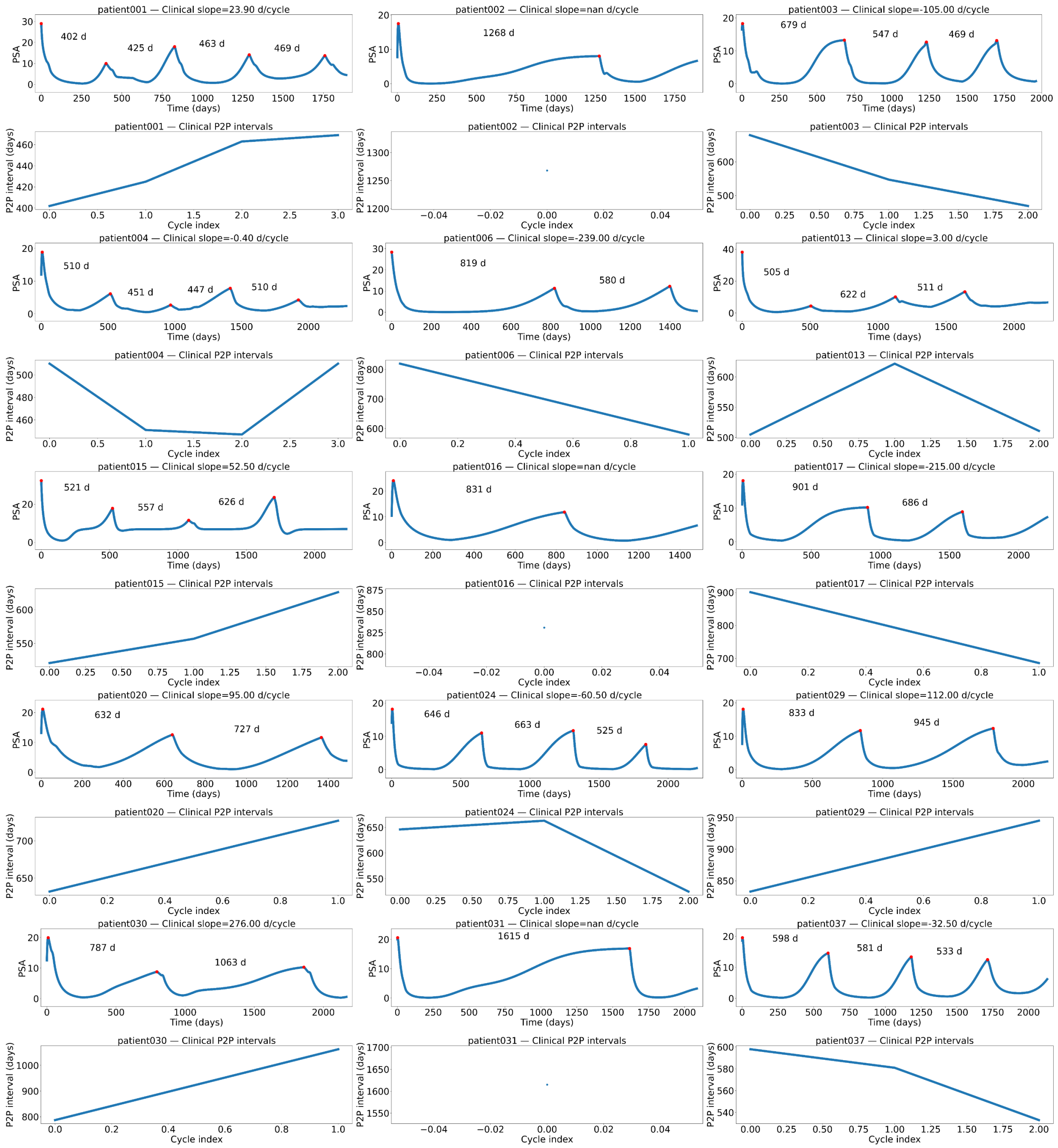


Figure S12: Cycle-wise response intervals under IADT (Part I). Top: PSA with cycle markers; bottom: P2P vs. cycle index with linear slope (days/cycle).

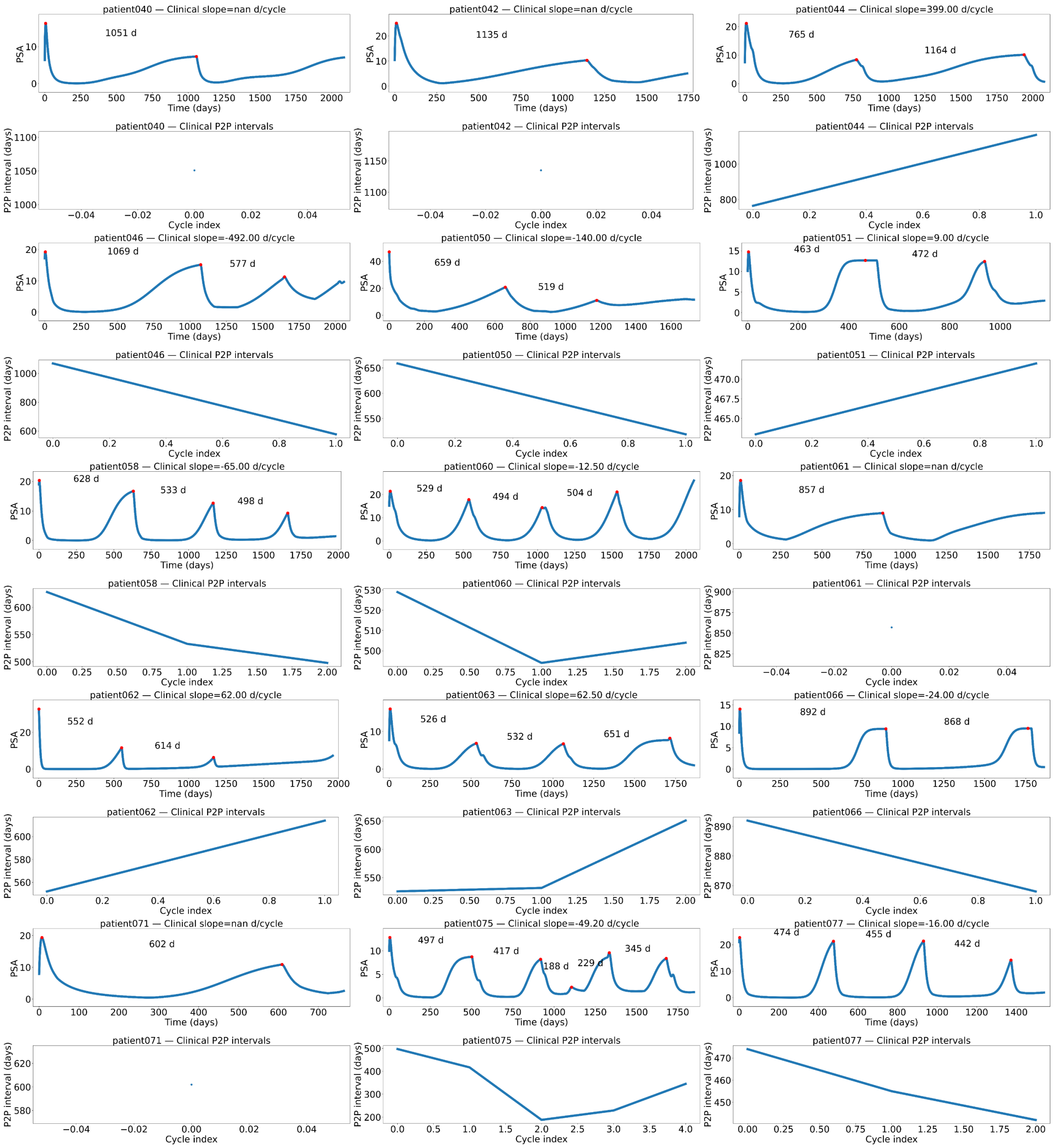


Figure S13: Cycle-wise response intervals under IADT (Part II).

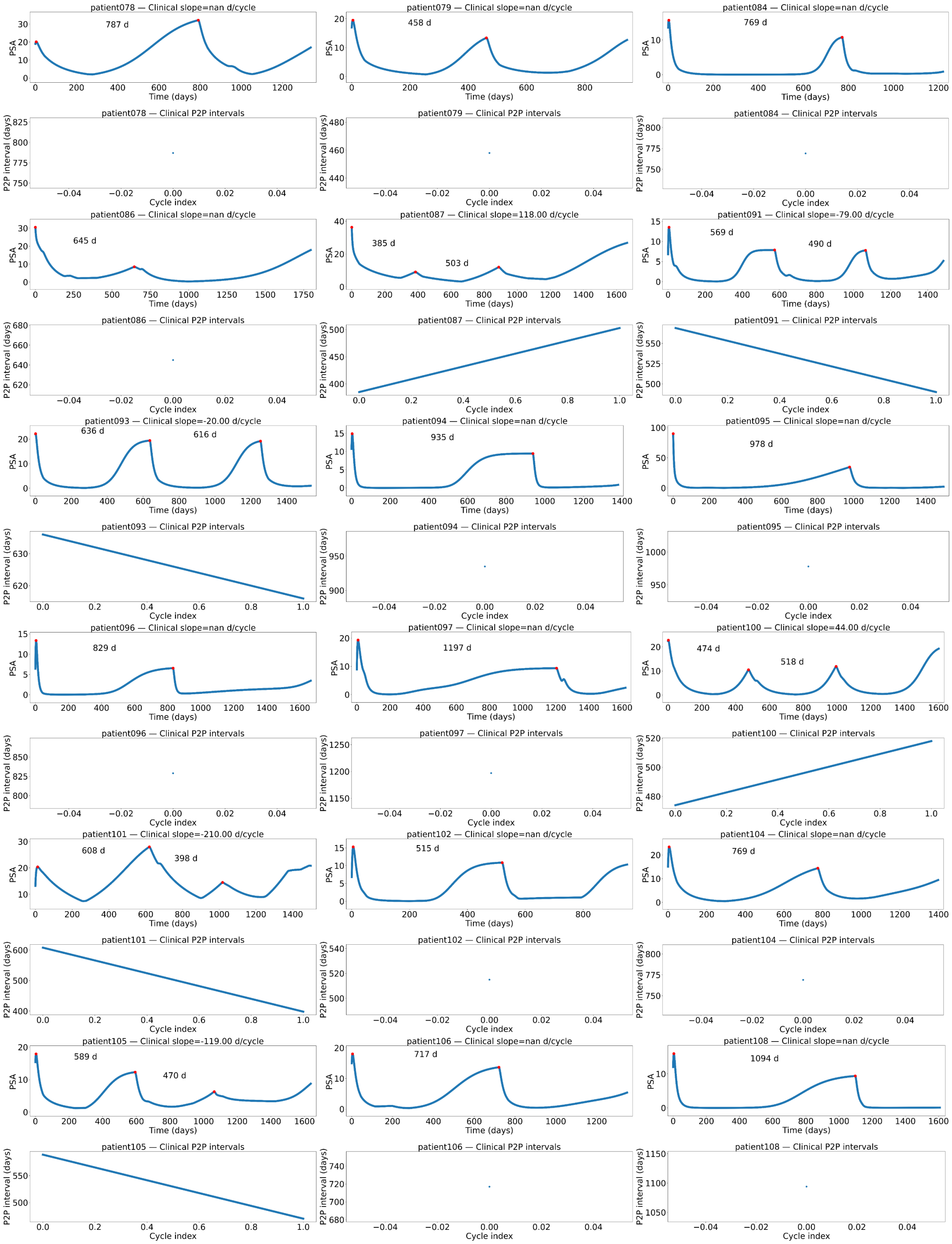


Figure S14: Cycle-wise response intervals under IADT (Part III).

### S8 Patients With vs. Without Cycle Shortening

Patients in the shortening-cycle group (Figure S15) exhibited progressively shorter treatment-response intervals across successive IADT cycles, reflecting gradual loss of tumor control and earlier onset of resistance. In contrast, patients in the non-shortening group (Figure S16) did not show a consistent contraction of cycle lengths; instead, their response intervals fluctuated across cycles, sometimes lengthening or showing local minima and maxima, indicative of more durable though variable responses. These two trajectories highlight distinct adaptive dynamics within the cohort, underscoring the clinical relevance of cycle shortening as a marker of tumor adaptation. Patients who experienced only a single treatment cycle could not be classified into either of these categories. Among the patients with progression, Patient012 was the only case showing cycle shortening, whereas all other patients with progression underwent just a single cycle of treatment and thus fell outside both the shortening and non-shortening groups.

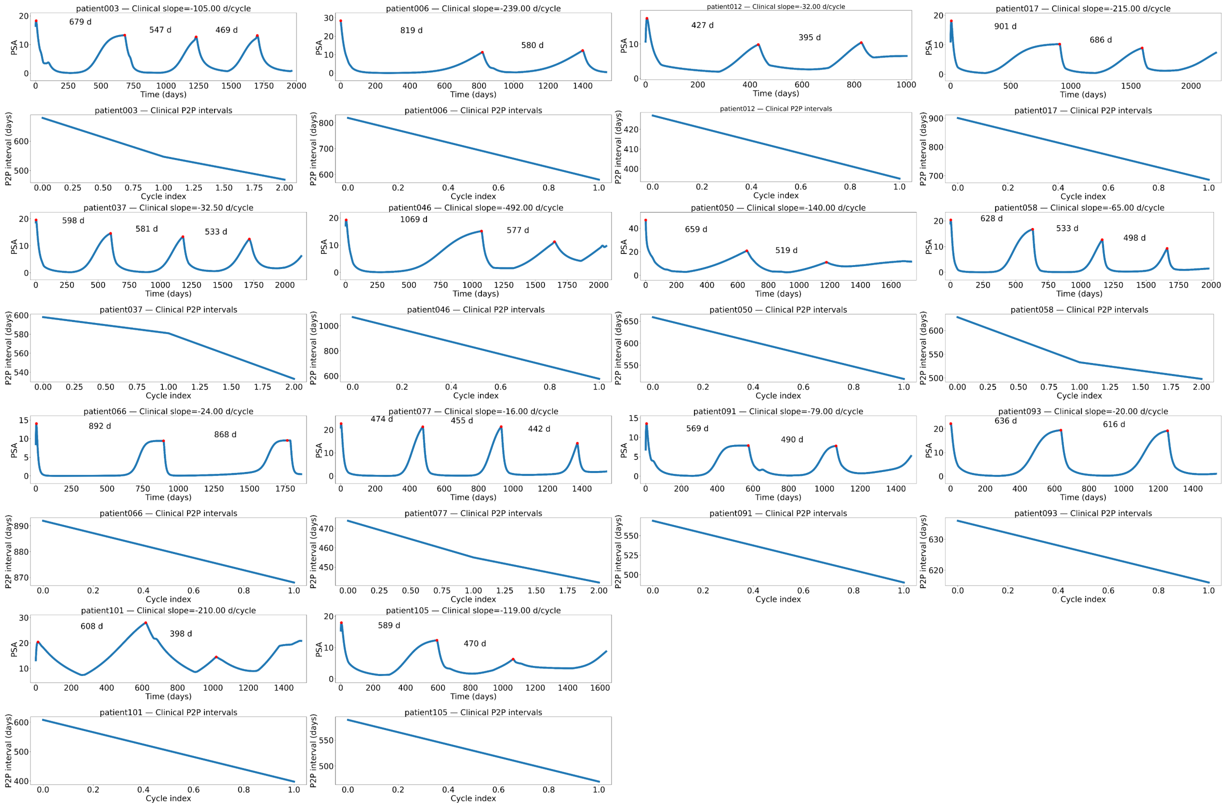


Figure S15: Shortening-cycle patients: progressively shorter response intervals. Patients in the shortening-cycle group exhibit progressively shorter treatment-response intervals across successive IADT cycles, reflecting gradual loss of tumor control and earlier onset of resistance.

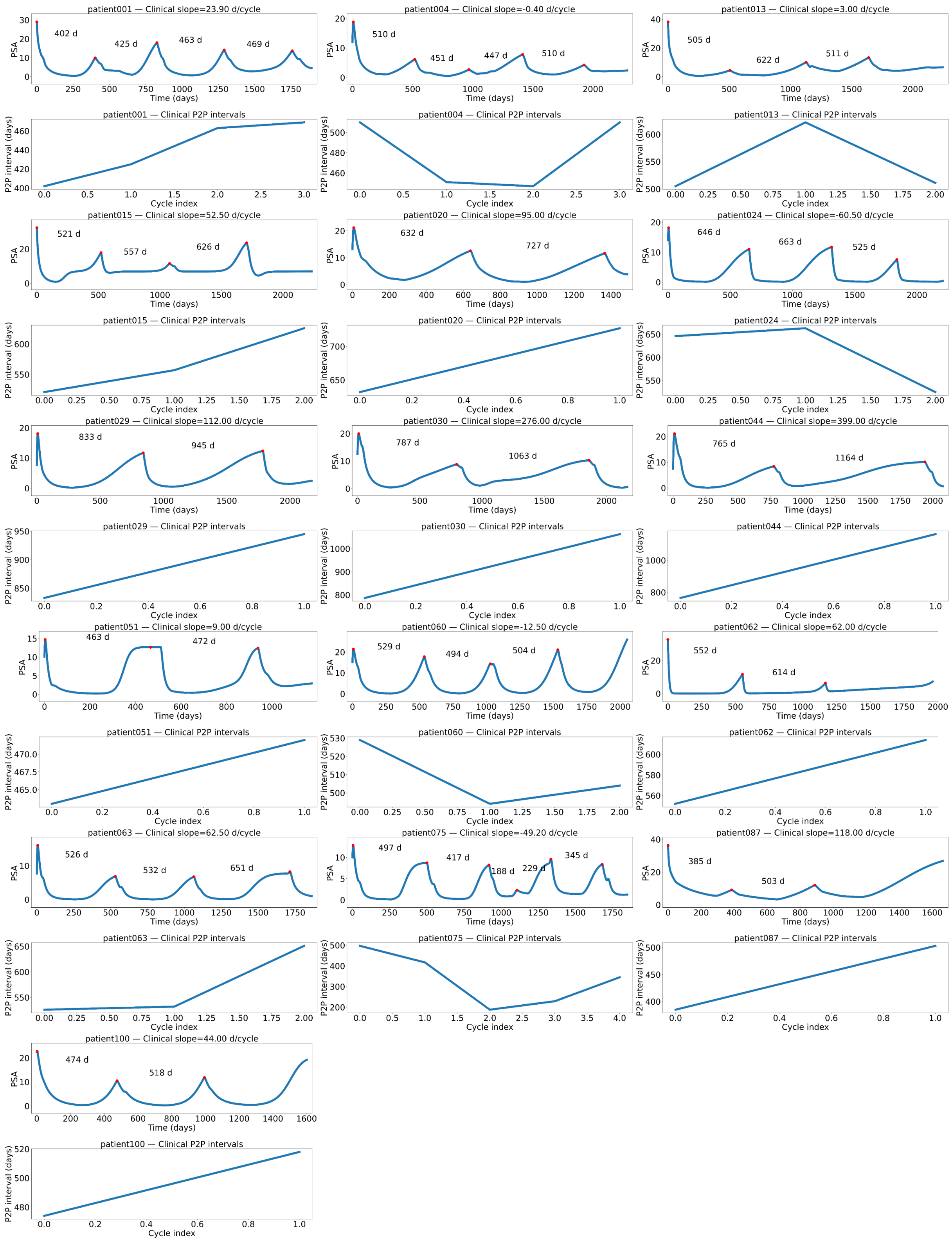


Figure S16: Non-shortening patients. Patients in the non-shortening group maintain relatively stable cycle lengths over time, indicative of more durable responses and delayed resistance.
